# FIKK1, a member of the FIKK kinase family, phosphorylates VAR2CSA and regulates adhesion of *Plasmodium falciparum*-infected erythrocytes to the placental receptor CSA

**DOI:** 10.64898/2026.03.23.713582

**Authors:** Benoît Gamain, Jean-Philippe Semblat, Heledd Eavis, Hugo Belda, Romain Hamelin, Clara-Eva Paquereau, Christian Doerig, Moritz Treeck, Dominique Dorin-Semblat

## Abstract

*Plasmodium falciparum* promotes the adhesion of infected erythrocytes (IEs) to host cells by extensively remodeling their surface. For this process, the parasite exports a large number of proteins to its host erythrocyte, including members of the *P. falciparum* Erythrocyte Membrane Protein-1 (PfEMP1) adhesin family and members of the FIKK family. Several FIKK have been shown to play a role in *P. falciparum* virulence, notably affecting IES cell surface remodeling, rigidity and cytoadhesion. VAR2CSA, a member of the PfEMP1 adhesin family, is associated with IES sequestration in the placenta and has been shown to be phosphorylated. In view of the previously described importance of VAR2CSA phosphorylation, we investigated the role of FIKK1. We show that FIKK1 is capable of phosphorylating VAR2CSA *in vitro*, and that this phosphorylation increases the binding of recombinant VAR2CSA to the placental receptor chondroitin sulphate A (CSA). In an inducible transgenic cell line expressing HA-tagged FIKK1, immunofluorescence assays indicate that the kinase localizes to punctuated foci within Maurer’s Cleft, similarly to VAR2CSA. Rapamycin-induced knock out of FIKK1 reduces IEs cytoadhesion to CSA, even though levels of VAR2CSA are not affected. In *vitro* phosphorylation assays show that FIKK1 can phosphorylate recombinant DBL1-3 domains on several residues, including S429 and T934, previously implicated in *in vitro* binding and IES cytoadhesion to CSA. Taken together, these data support a model whereby FIKK1 contributes to placental malaria virulence through IEs sequestration mediated by VAR2CSA phosphorylation. Having no orthologs in mammals, this orphan kinase therefore represents an attractive target for the development of drugs against placental malaria.

**Author summary:** Sequestration of *Plasmodium falciparum* infected erythrocytes (IEs) in the placenta is a hallmark of placental malaria and a major driver of adverse maternofetal outcomes. This process is mediated by VAR2CSA, a member of the *P. falciparum* Erythrocyte Membrane Protein (PfEMP1) family. VAR2CSA binds to chondroitin sulfate A (CSA) on the placental syncytium, facilitating IEs sequestration. In a previous study, we demonstrated that endogenous VAR2CSA is phosphorylated and identified specific phosphosites important for its cytoadhesive function. To elucidate the molecular mechanisms underlying these post-translational modifications, we examined the role of the *P. falciparum* FIKK1 kinase in VAR2CSA phosphorylation and its impact on IEs adhesion. Using a FIKK1::HA conditional knockout transgenic line, we found that FIKK1 deletion impairs IEs cytoadhesion, likely due to altered VAR2CSA phosphorylation. Importantly, both endogenous and recombinant FIKK1 interact with and phosphorylate the extracellular region of VAR2CSA. Furthermore, recombinant FIKK1 also enhances VAR2CSA binding to CSA *in vitro* and phosphorylates a residue previously identified as important for CSA binding and IEs adhesion. Collectively, these findings highlight a pivotal role for FIKK1 in placental adhesion and underscore the potential of targeting this kinase family for interventions against placental malaria.

## Introduction

Malaria caused by *Plasmodium falciparum* remains one of the world’s most severe infectious diseases and a global health priority. According to the WHO’s latest *World malaria report*, there were an estimated 263 million cases and 597,000 deaths caused by malaria worldwide in 2023 [1]. Although, globally, the prevalence of malaria has shown a gradual decline over the years, malaria continues to be a major public health challenge [2]. The disease mostly affects young children and pregnant women in endemic areas [3]. The severity of the disease is mainly associated with the cytoadhesion of infected erythrocytes (IEs) to various receptors expressed on the surface of host cells [4]. The polymorphic *var* gene family, encoding an array of antigenic variant molecules called *Plasmodium falciparum* Erythrocyte Membrane protein 1 (PfEMP1), plays a central role in IEs cytoadhesion, and hence is a major determinant of virulence [5]. PfEMP1 is exported to the IEs cytoplasm and anchored into IEs plasma membrane protrusions called knobs. These adhesins mediate IEs binding to a variety of receptors on the vascular endothelium in various organs such as the brain or the placenta. VAR2CSA is the PfEMP1 responsible of IEs sequestration in the placenta, through binding to its specific ligand Chondroitin Sulfate A (CSA), which is expressed on the surface of syncytiotrophoblasts [6, 7]. IEs cytoadhesion to the placental layer leads to severe outcomes for both the mother and the fœtus, resulting in low birth weights or stillbirth [8].

Phosphorylation by kinases is a key biochemical process involved in the regulation of cell adhesion [9], by turning “on” or “off” the interaction between the adhesin molecules and their ligand. Our previous study strongly suggests that the phosphorylation of the extracellular region of the antigenic variant VAR2CSA enhances adhesive properties of IEs to CSA, and crucial phosphosites mediating this regulation have been identified [10].

Identification of novel drugs with untapped targets are urgently needed to face the emergence and propagation of drug-resistant parasites. Protein kinases have already been successfully targeted to treat numerous diseases [11] [12]. Protein kinases play a pivotal role in malaria by controlling *P. falciparum* multiplication and pathogenicity; therefore, inhibiting these enzymes represents a potential avenue for the prevention or treatment of *P. falciparum* malaria [13]. The FIKK kinase family, expanded to 20 members in *P. falciparum*, is named after the presence of a conserved (F)Phenylalanine-(I) Isoleucine (K) Lysine (K) Lysine amino acid motif [14]. This lineage-specific kinase family is considered as a potential and promising target. FIKK kinases are exclusively found in Apicomplexa (a taxon of parasitic Alveolates containing a specific apical organelle important for host invasion). These kinases are absent in humans and other eukaryotes, and most Apicomplexa possess only a single *fikk* gene, making members of this gene family potential attractive biomarkers for species-specific diagnostics [15] [16]. In addition to a conserved C-terminal catalytic kinase-like domain, all FIKKs possess a variable N terminal extension. The essentiality of some of these enzymes for parasite blood stage growth, and their location in different IES compartments, suggest that FIKKs have various, non-redundant roles [17]. Most FIKKs contain a conserved export motif termed *<u>P</u>lasmodium* <u>ex</u>port <u>el</u>ement (PEXEL) mediating their export to the host RBCs [18]. This protein export system involves parasite-derived membranous structures found in the host erythrocyte’s cytoplasm, called Maurer’s Clefts (MCs); these constitute an important intermediate trafficking compartment from the parasite to the host cell membrane [19] [20].

Specific antibodies against various members of the FIKK family reveal the presence of some FIKKs in the IEs cytoplasm, displaying a punctuated pattern similar to that shown by MC resident proteins [18]. Phosphorylation of cytoskeletal erythrocyte proteins such as dematin by FIKK4.1 [21], or spectrin, ankyrin and Band-3 by FIKK9.1 [22], suggest a key role for these enzymes in host cell remodeling, a process critical for *P. falciparum* infection. Furthermore, targeted disruption of the *fikk4*.*2* gene dramatically alters remodeling of structures under the plasma membrane of the host erythrocyte, affecting its rigidity and adhesive properties to endothelial receptors such as CD36 [23].

Combined approaches based on inducible conditional knock out of FIKK genes associated to global quantitative phosphoproteomics [24] brought new insights into: i) the location of all FIKKs predicted to be exported, ii) the identification of a unique phosphorylation fingerprint for each kinase FIKK family member, iii) the complex network that FIKKs belong to and iv) the identification and position of the phosphosites in several parasite and host cytoskeletal proteins phosphorylated by FIKK4.1. Additionally, using combined structural analysis and AlphaFold 2 predictions, residues that determine target motif specificity for some FIKKs were identified [25]. Adducin, protein 4.1 and KHARP, previously identified as interactors of FIKK4.1 and FIKK4.2 [26], are phosphorylated by both kinases on non-overlapping residues, in line with *in silico* predictions. Furthermore, adducin was shown to be phosphorylated specifically by FIKK1 on S726 [24]. Overall, available data point to a role of FIKKs in IEs rigidity and cytoadhesion. In the present study, we assessed the putative role of FIKK1 in phosphorylating VAR2CSA and regulating IEs cytoadhesion to the placental receptor CSA, using a NF54::DiCre transgenic cell line expressing a FIKK1 fused to a C-terminal HA tag [27] [24]. Notably, we show that HA-tagged FIKK1 displays a similar staining to VAR2CSA as punctuated foci in Maurer’s Clefts and that FIKK1 can phosphorylate *in vitro* recombinant VAR2CSA on several residues, notably within sites previously implicated in *in vitro* binding and IEs cytoadhesion to CSA. Deletion of the kinase impairs the ability of VAR2CSA-expressing IEs to cytoadhere to CSA. Taken together, our data identify FIKK1 to play a role in regulating *P. falciparum* pathogenicity.

## Results

### VAR2CSA is expressed in the NF54 FIKK1::HA transgenic clone F5 and localizes with FIKK1 at Maurer’s cleft structures

The previously described *fikk1* gene conditional knockout (cKO) line, expressing FIKK1 fused at its C-terminal extremity to a triple hemagglutinin HA-tag [24], was used to assess the role of the kinase in modulating IEs CSA-binding. To ascertain homogeneity of the parasites, the *fikk1* cKO transgenic cell line was recloned by limiting dilution and the F5 clone was characterized by flow cytometry for VAR2CSA expression. FACS analysis revealed that 90% of F5 clone IEs express VAR2CSA on their surface **(Fig 1A)**. Western Blot of IEs total extracts with a purified anti-VAR2CSA goat antibody confirms the presence of VAR2CSA at the expected molecular weight **(Fig 1B upper right panel)** with a slightly higher molecular weight than the recombinant VAR2CSA rDBL1-6 positive control (lane 1) as expected. In the same extracts, an anti-HA western blot confirms the presence of HA-tagged FIKK1 at the expected size (∼82kDa) in the non-induced cKO FIKK1::HA transgenic line, but not in the NF54 WT strain **(Fig 1B bottom right panel)**.

**Fig 1.**
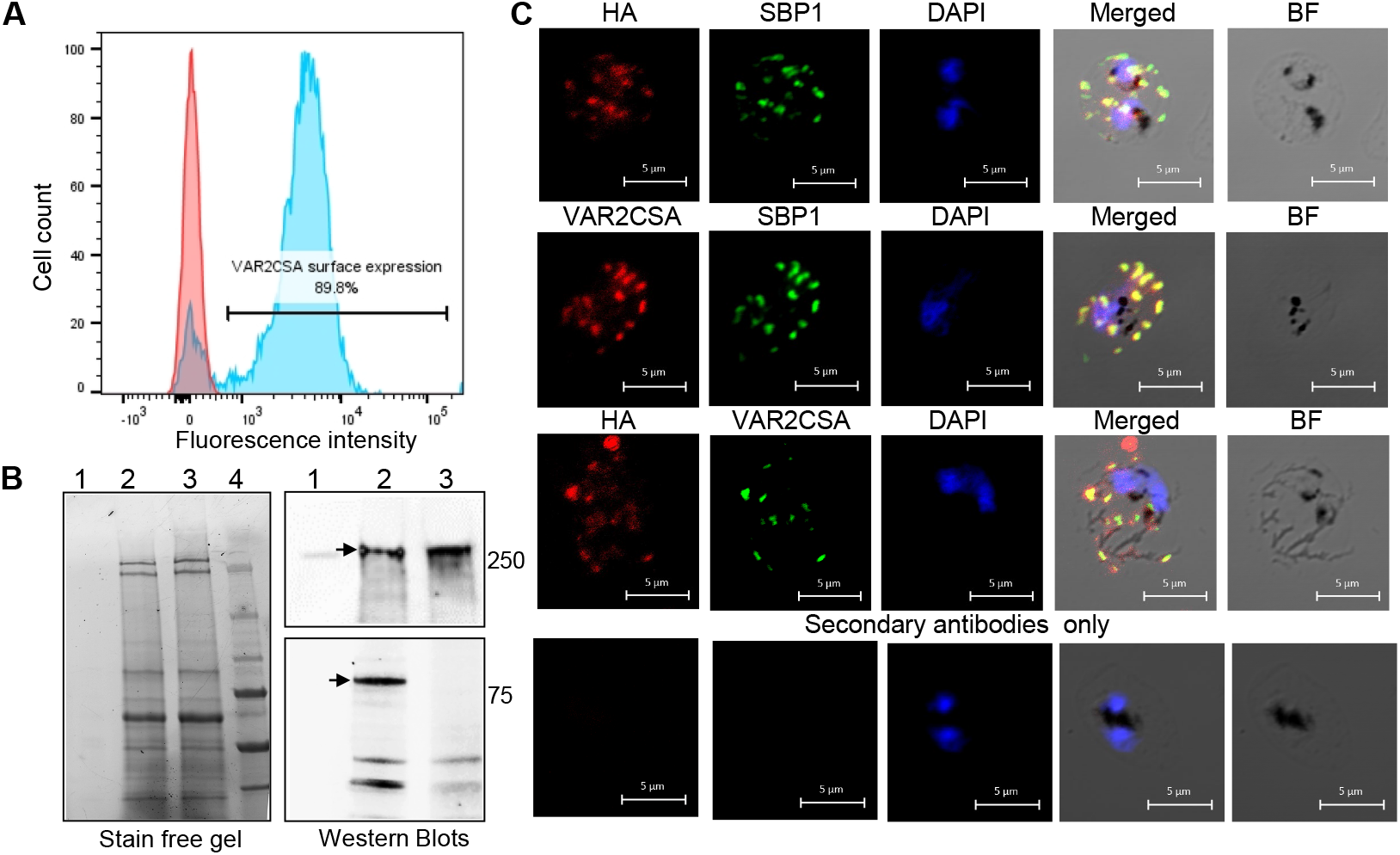
Expression and localization of PfEMP1 VAR2CSA and FIKK1 localization in FIKK1::HA conditional knock out transgenic line. **A.** Flow cytometry analysis of cKO FIKK1::HA: transgenic cell line using specific anti-VAR2CSA antibodies. **B**. AntiVAR2CSA (upper panel) and anti-HA Western Blot (bottom panel) were performed on total protein lysates of the cKO FIKK1::HA cell line. The corresponding stain free gel is shown on left panel to visualize the amount of each lysate loaded on the gel. Lane 1: recombinant extracellular region of VAR2CSA (rVAR2CSA DBL1-6); lane 2: NF54 cKO FIKK1::HA lysate; lane 3: NF54VAR2CSA WT lysate; Lane 4: Molecular weight. **C**. Immunofluorescence assays of erythrocytes infected with the FIKK1::HA cell line using antibodies against the C-terminal HA-tag. For subcellular localization of FIKK1 and VAR2CSA, fixed and permeabilized smears of mature stages were dual labelled *either* with anti-HA / PfSBP1 (Maurer’s Cleft), *anti-*VAR2CSA / PfBSP1 *or with anti HA / VAR2CSA* antibody pairs. *Scale bar is indicated*

The HA tag allowed us to visualize the subcellular localization of FIKK1 on fixed and permeabilized IEs by immunofluorescence assays (IFAs) using an anti-HA antibody. IFAs revealed punctate staining outside of the parasite and within the IEs cytoplasm, reminiscent of the previously described Maurer’s clefts staining pattern [19]. To determine whether the observed foci in the IEs cytoplasm was indeed associated with the latter compartment, we examined their co-localization with *P. falciparum* Skeleton Binding Protein 1 (PfSBP1), a known resident protein of the MCs. FIKK1-HA displays a highly similar staining pattern to that of PfSBP1, consistent with the presence of the protein kinase in MCs **(Fig 1C)**.

IFAs with specific anti-VAR2CSA antibodies showed that VAR2CSA displays a similar staining pattern to that of PfSBP1, confirming recent data [28] **(Fig 1C)**. Co-labelling of FIKK1-HA and VAR2CSA shows a punctuated pattern reminiscent of MCs staining, suggesting that both proteins are located in the same intermediate trafficking compartment. This pattern was not observed in slides labelled with control isotype antibody **(Fig 1C)**.

### VAR2CSA interacts with FIKK1

The VAR2CSA-FIKK1 co-staining pattern in the MCs led us to hypothesize a putative interaction between VAR2CSA and FIKK1. To verify this hypothesis, we performed an immunoprecipitation of FIKK1-HA with an anti-HA mouse monoclonal antibody on both Triton-soluble (mostly cytosolic and some solubilized cytoskeleton-associated proteins) and Triton-insoluble (mostly solubilized membrane and cytoskeleton-associated proteins) fractions obtained from IEs. These fractions will be respectively called soluble and membrane fractions. Control immunoprecipitations on both fractions were also performed with a mouse isotype antibody. Immunoprecipitated material was analyzed by western blot with an anti-HA and a specific anti-VAR2CSA goat antibody after migration on a stain-free gel (**Fig 2 A)**. The kinase is present in both fractions (blue arrow) but predominantly in the soluble fraction **(Fig 2A, right lower panel, lanes 5 and 6)**. Interestingly, anti-VAR2CSA antibodies revealed a faint band (indicated with a red arrow) corresponding to the size of VAR2CSA in the HA immunoprecipitated material from the membrane fraction, suggesting that FIKK1 interacts with VAR2CSA in this fraction **(Fig 2A, right upper panel, lane 5)**. This band was also present in the membrane lysates starting material **(Fig 2A, right upper panel, lane 1)**. No band was observed in the control immunoprecipitations performed with the isotype antibody.

**Fig 2.**
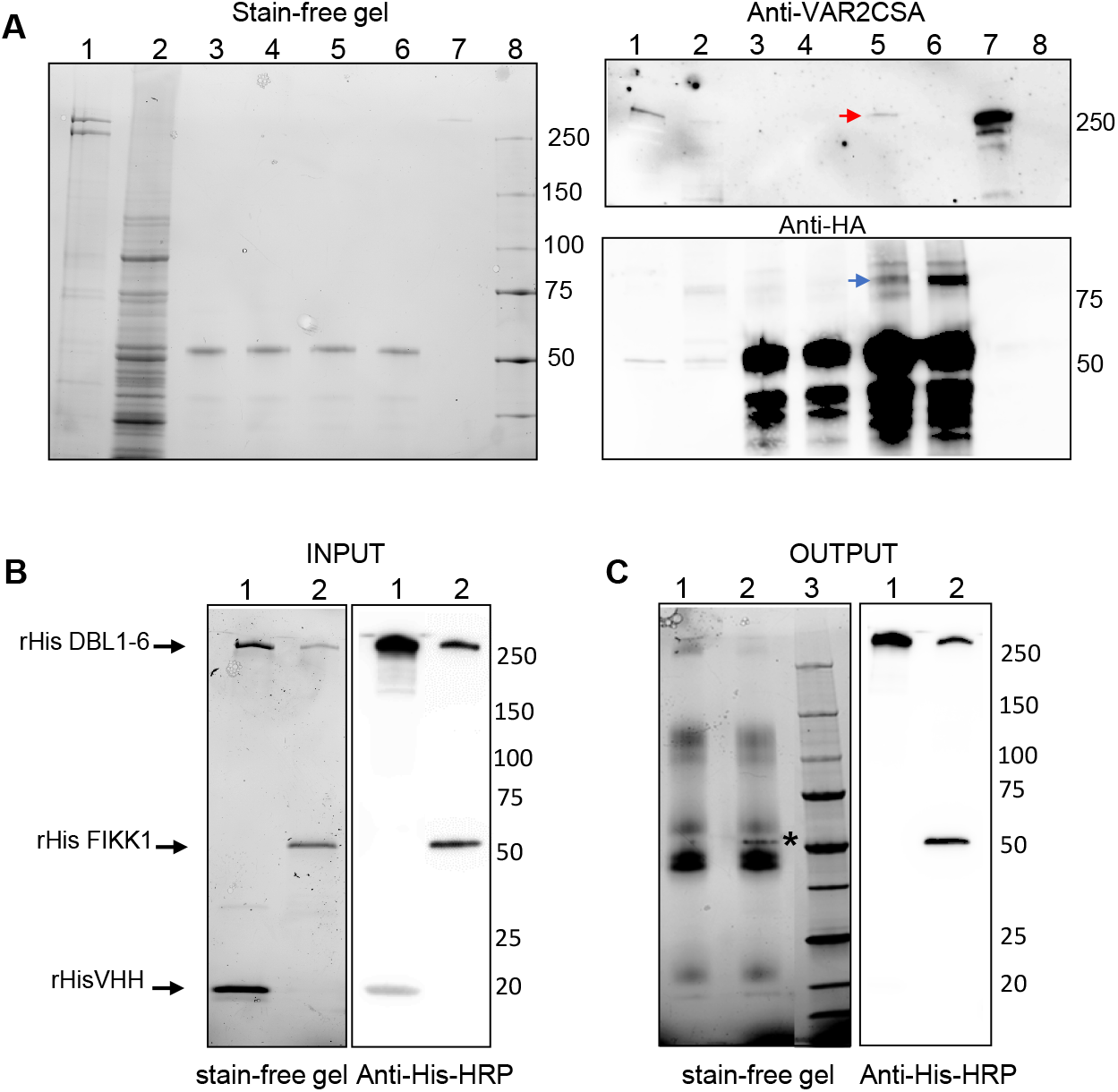
Native VAR2CSA co-immunoprecipitates with FIKK1::HA and rFIKK1 kinase domain interacts with rVAR2CSA DBL 1-6. **A**. Immunoprecipitation was performed on membrane (Triton insoluble) and soluble (Triton soluble) protein extracts from transgenic parasites expressing HA-tagged FIKK1 using either a mouse anti-HA or a mouse isotype control. Stain-free gel (left panel). Lane 1: Input membrane protein lysate; lane 2: input soluble protein lysate; lane 3: immunoprecipitation with an isotype antibody on membrane protein extracts; lane 4: immunoprecipitation with isotype antibody on soluble protein extracts; lane 5: immunoprecipitation with anti-HA on membrane protein extracts; lane 6: immunoprecipitation with anti-HA on soluble protein extracts. lane 7: recombinant VAR2CSA DBL1-6. lane 8: Molecular weight. Western blot analysis using anti-VAR2CSA antibodies (right upper panel). Immunoprecipitated VAR2CSA is shown with a red arrow. Western blot using a mouse anti-HA antibody (right lower panel). Immunoprecipitated FIKK1::HA is indicated with a blue arrow. **B**. Recombinant His-VHH or recombinant His-FIKK1 kinase domain (INPUT) were incubated with His-rDBL1-6 and were detected either by stain free gel (left panel) or by Western blot (right panel). Lane 1: rDBL1-6 + rVHH (input); lane 2: rDBL1-6 + rFIKK1 (input). **C**. Recombinant DBL1-6 was immunoprecipitated using specific antibodies and any bound proteins to rDBL1-6 (OUTPUT) were detected by stain-free SDS gel (left panel) prior western blotting (right panel). Lane 1: rDBL1-6 + rVHH (bound fraction); lane 2: rDBL1-6 + rFIKK1 (bound fraction). Lane 3: molecular weight. Recombinant proteins are indicated with arrows and rFIKK1 bound to rDBL1-6 is indicated with a * on the stain free gel.

The above results suggest that VAR2CSA can directly or indirectly interact with FIKK1. To further address this question, we performed *in vitro* binding assays with purified His-tagged recombinant proteins. Recombinant VAR2CSA extracellular region rDBL1-6 was mixed either with recombinant FIKK1 kinase domain or with an irrelevant camelid <u>V</u>ariable domain of <u>H</u>eavy chain of <u>H</u>eavy-chain only antibody from Camelids (VHH) antibody as a negative control. All proteins from the starting material samples (INPUT) are visualized by stain-free gel **(Fig 2B, left panel)** and by western blot using an anti-His antibody **(Fig 2B, right panel)**. Rabbit anti-VAR2CSA antibodies bound to dynabeads were added to immunoprecipitate recombinant DBL1-6 and any bound protein. Stain-free gel of the bound material (OUTPUT) shows a band around 50kDa (indicated as a *) corresponding to the FIKK1 kinase domain (KD) just above the IgG band but no band of the size of VHH **(Fig 2C, left panel)**. An anti-His western blot confirms the presence of the recombinant FIKK1 and not the irrelevant VHH in the VAR2CSA immunoprecipitated materials **(Fig 2C, right panel)**. Additionally, ELISA binding assays were carried out using rVAR2CSA DBL1-6 added in a plate precoated with rFIKK1 KD, or with the irrelevant VHH as a negative control or PBS. A monoclonal anti-VAR2CSA antibody and an anti-mouse HRP conjugated secondary antibody were used to detect bound rDBL1-6. Results are displayed on a histogram and show almost a 10-fold increased binding between rVAR2CSA DBL1-6 and rFIKK1KD compared to rDBL1-6 and rVHH **(S1 Fig)**. This strongly suggests that FIKK1 and VAR2CSA interact *in vitro*.

### VAR2CSA expression is unchanged after Rapamycin-mediated deletion of FIKK1

Having shown that VAR2CSA is the major PfEMP1 expressed on the surface of FIKK1::HA clone F5 IEs and furthermore localizes and interacts with FIKK1, we then investigated a possible role for FIKK1 in regulating VAR2CSA surface expression and on IES adhesion to CSA. We conditionally deleted FIKK1 by treating the FIKK1::HA cKO strain F5 clone (synchronized at the ring stage) with rapamycin (or the DMSO vehicle as a control). We first verified that rapamycin treatment leads to the deletion of the *fikk1* locus 72 hours post induction. As expected, efficient loss of the HA tagged gene was verified by PCR performed directly on the culture and on purified genomic DNA Specific pairs of primers confirmed the expected size for 5’ and 3’recombination events and locus excision after rapamycin treatment **(S2 Fig**). To ensure that rapamycin treatment leads to the absence of the HA-tagged protein kinase, we performed an IFA using anti-HA antibody. Analysis of fixed and permeabilized asexual parasites reveals a fluorescent staining in several DMSO treated parasites but not in RAPA-treated parasites **(Fig 3A)**. Deletion of the gene locus was confirmed by anti-HA western blot **(Fig 3B right panel)** on total protein lysates: a ∼ 80kDa band present in the FIKK1::HA lysates disappears in RAPA-treated parasites. The left panel is a stain free gel prior blotting to verify the equivalent amount of both lysates loaded on the gel.

**Fig 3.**
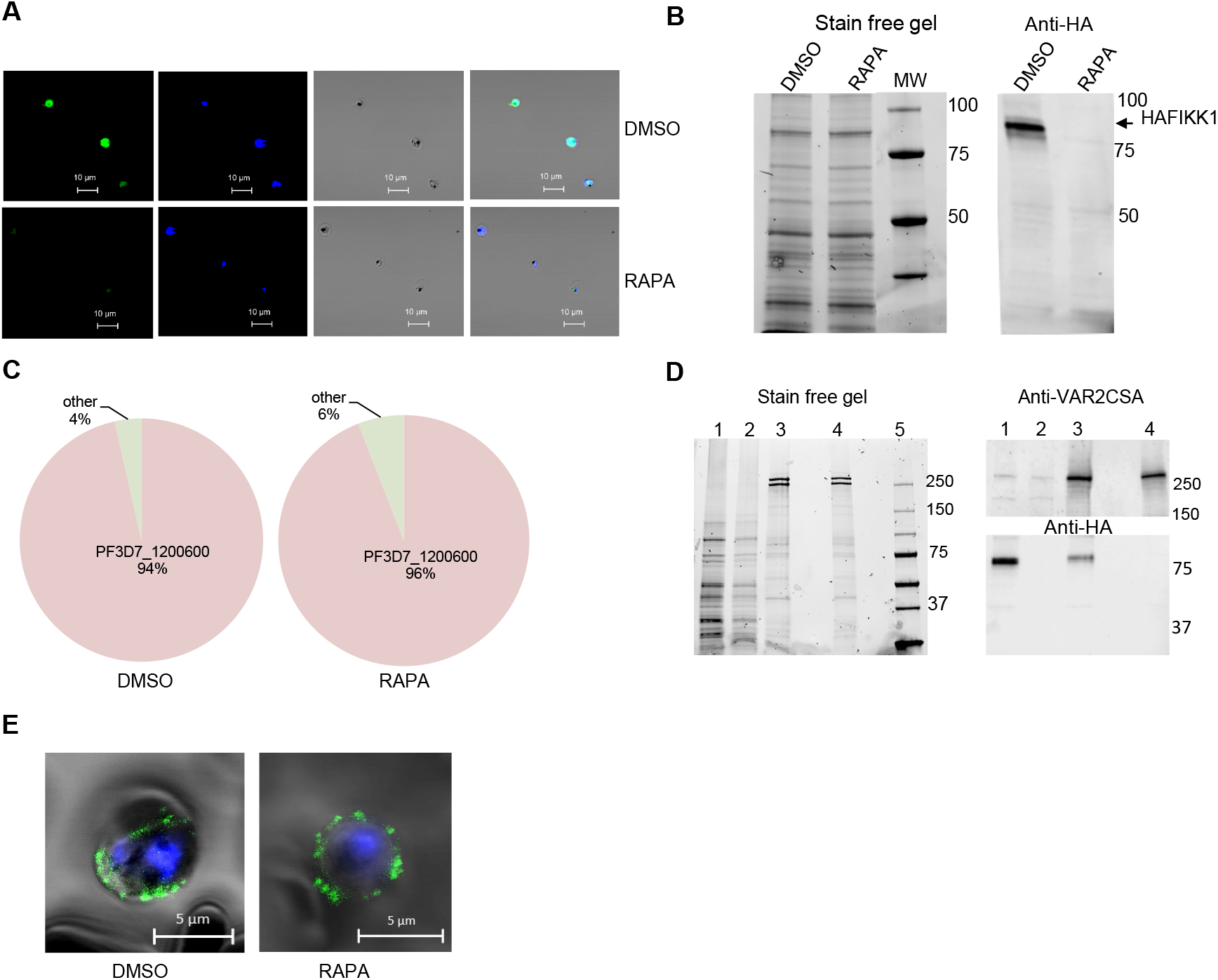
Rapamycin treatment induces FIKK1 KO and does not affect VAR2CSA expression. **A.** Immunofluorescence labelling of several DMSO and RAPA-treated asexual parasites using anti-HA antibody followed by a secondary Alexa Fluor 488-labelled antibody. **B.** *Anti-HA* Western blot (right panel). FIKK1 deletion is shown upon RAPA treatment. A stain free gel confirms equal loading of both total lysates (left panel). **C.** VAR2CSA transcription profile of DMSO and RAPA-treated asexual parasites. Transcriptional levels of each *var* genes were normalized with the housekeeping gene, seryl-tRNA transferase. **D.** Western blot with anti-VAR2CSA antibodies performed on soluble and membrane fractions of DMSO and RAPA-treated asexual parasites (upper right panel). Anti-HA western blot was assessed on the same protein lysates (lower right panel). Profile of soluble and membrane fractions was visualized on a stain free gel prior blotting (left panel). Lane 1: DMSO soluble lysate; lane 2: RAPA soluble lysate; lane 3: DMSO membrane lysate; lane 4: RAPA membrane lysate; lane 5: molecular weight. **E**. Immunofluorescence assays on live mature stages RAPA-treated cells and control DMSO-treated cells using anti-VAR2CSA antibodies. Scale bar is indicated.

Quantitative RT-PCR of *var* genes performed on both DMSO and RAPA treated -treated cell lines indicates the same percentage of *var2csa* transcripts (∼95%) suggesting that deletion of FIKK1 does not lead to a switch in *var* gene expression **(Fig 3C)**. Furthermore, no difference was observed in the fractionation of endogenous PfEMP1 VAR2CSA protein visualized by western blot with an anti-VAR2CSA performed on soluble and membrane lysates of DMSO **(Fig 3D lane 1 and 3 upper panel)** and RAPA-treated parasites (**Fig 3D lane 2 and 4 upper panel**). A Western blot probed with anti-HA antibodies of the same lysates showed the presence of the tagged kinase in soluble and membrane lysates in the DMSO treated lysates and its absence from extracts from knocked-out parasites, as expected **(Fig 3D lane 2 and 4 lower panel)**

### Rapamycin-mediated truncation of FIKK1 does not change VAR2CSA surface expression but impairs cytoadhesion

A previous study had established that the levels of PfEMP1 on the surface of IEs are not altered upon deletion of any FIKK except FIKK4.1 [26]. In line with these data, no change was observed in the level of VAR2CSA surface expression of the clone F5 used in this study, confirming that FIKK1 absence does not impair trafficking of the adhesin **(S3 Fig)**. Surface labelling IFA with specific antibodies displayed peripherical punctuated dots, which most likely are the knobs present at the IEs surface. No difference in the staining pattern was observed between DMSO- and RAPA-treated cell lines **(Fig 3E)**.

DMSO- or RAPA-treated parasites were then assessed for CSA cytoadhesion phenotype using static adhesion assays. Interestingly, despite similar VAR2CSA surface expression **(S3 Fig)**, a significant decrease of the number of bound IEs was observed after Rapamycin treatment, when compared to the DMSO-treated cell line using static binding assays on immobilized CSA (average of 60% reduction; **p=0.002; n=3) **(Fig 4 A and B)**. This inhibition was further confirmed, (average of 50% reduction; ****p=0.0001; n=3), using a 96-well plate CSA binding assay **(Fig 4C)**. In order to rule out that the observed static adhesion phenotype is due to an inhibitory effect of the Rapamycin treatment, these experiments were performed on the NF54 WT VAR2CSA-expressing parental line. No change in VAR2CSA surface expression and no adhesion defect was observed upon RAPA-treatment for the NF54 parental line **(S4 Fig)**, confirming that the observed phenotype is due to the absence of the kinase in the RAPA-induced FIKK1 cKO line. We assessed IEs cytoadhesion also in flow conditions that mimic the circulatory system in placenta. DMSO- and RAPA-treated. The analysis of the ratio before and after a 4.5mL/h flow rate indicates a reduction in the binding of RAPA-treated parasites relative to DMSO-treated (although not significant). A significant reduction was observed by increasing the flow rate to 150mL/h **(Fig 4D)**. Therefore, the above data concur to indicate that FIKK1 regulates VAR2CSA-mediated IEs cytoadhesion without altering the adhesin expression levels.

**Fig 4.**
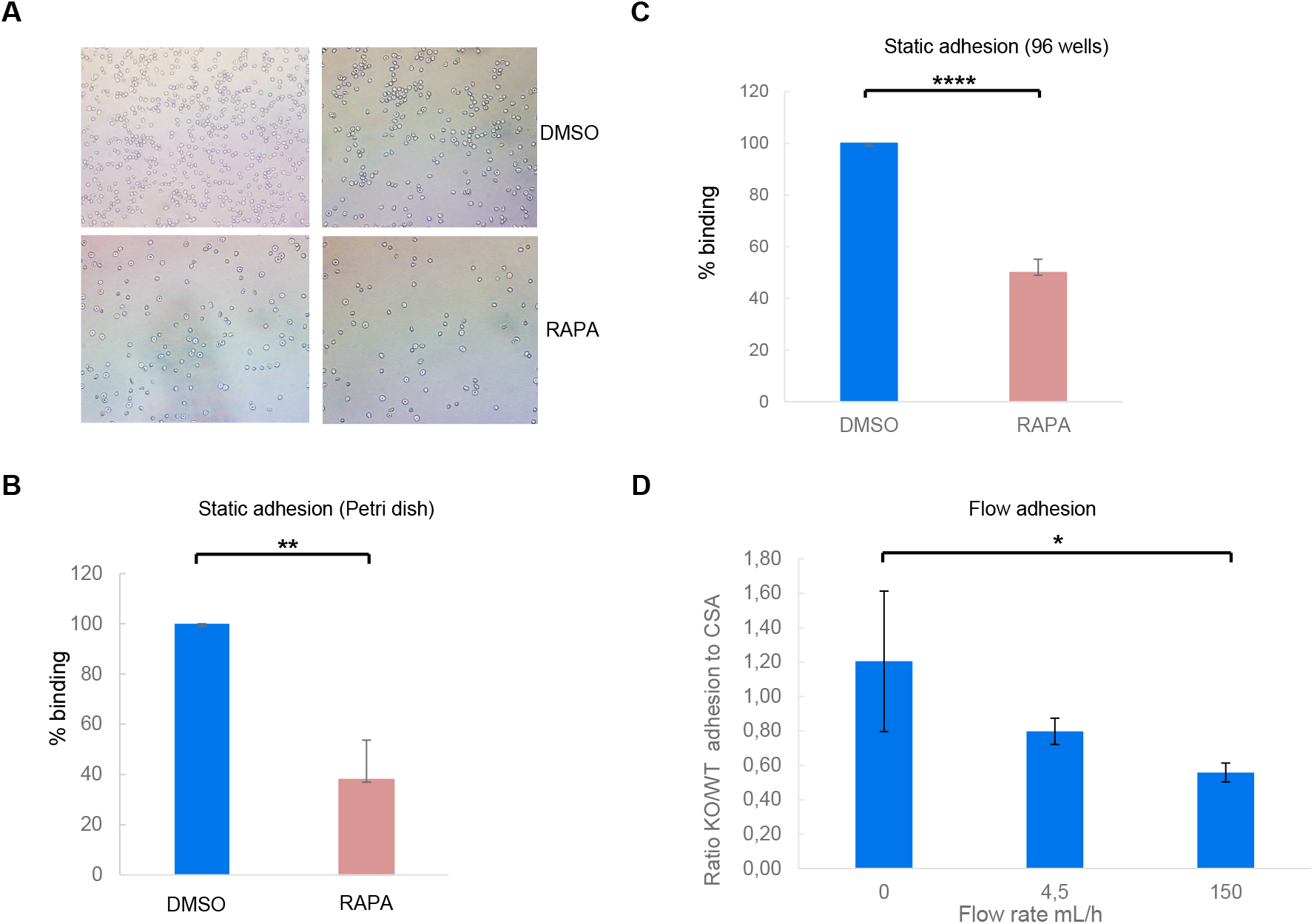
Deletion of FIKK1 after rapamycin induction affect IEs cytoadhesion. **A and B**. Static adhesion assay on plastic coated CSA. **A**. Typical pictures of bound DMSO and RAPA-treated IEs conditions. **B**. Bound IEs were counted in 5 random fields. Results were expressed as percentage of binding (DMSO-treated cells correspond to 100% binding). Mean and standard deviation are indicated. (Statistics: n=3, Unpaired t test **p=0.002). **C**. Static adhesion assay in 96 wells plate. CSA-binding inhibition was assessed by relative quantification of DMSO and RAPA-treated IEs remaining bound to the plate surface after washes The OD _650nm_ measures were obtained in triplicate; Means were used for calculation; Experimental data were analyzed using GraphPad Prism software. (Statistics: n= 3, ****p<0.0001 Unpaired t test). **D**. Flow adhesion of RAPA/DMSO treated IES over CSA-coated slides at 4.5 mL/hr and 150 mL/hr flow rates; (Statistics: n= 6, *p=0.02 Paired t test).

### FIKK1 phosphorylates VAR2CSA

We then investigated the capacity of the endogenous FIKK1 to phosphorylate the extracellular region of VAR2CSA. To purify endogenous FIKK1, an immunoprecipitation with the mouse monoclonal anti-HA antibody was performed on total lysates from DMSO- and Rapamycin-treated transgenic lines. An aliquot of the immuno-conjugated material was used to perform a western blot with an anti-HA antibody. A band of approximatively 80kDa was detected in the DMSO-treated parasites and absent in the RAPA-treated cells, confirming the deletion of the kinase **(Fig 5 A)**. Immuno-conjugated material was used as well as a source of kinase to phosphorylate the recombinant extracellular domain of VAR2CSA with ATPγ32P. A strong radiolabeled band corresponding to the size of the recombinant extracellular region was present on the autoradiogram with the DMSO-treated material **(Fig 5B, right panel, lane 2)** whereas a weak band was observed with the RAPA-treated material **(Fig 5B, right panel, lane 1)**. This result indicates that the endogenous FIKK1 is able to phosphorylate rVAR2CSA extracellular domain (although an indirect effect involving another kinase cannot be formally ruled out, see below). Interestingly a weak band of around 80 kDa was visualized in the autoradiogram in the lane corresponding to the DMSO control (lane 2) but not in immunoprecipitated material from the RAPA treated cells (lane 1). This band may arise from FIKK1-HA autophosphorylation. Interestingly, we observed a radiolabeled band with a high molecular weight (>300kDa) corresponding to the size of full-length VAR2CSA (indicated with a *) just above the labelled recombinant VAR2CSA extracellular region

**Fig 5.**
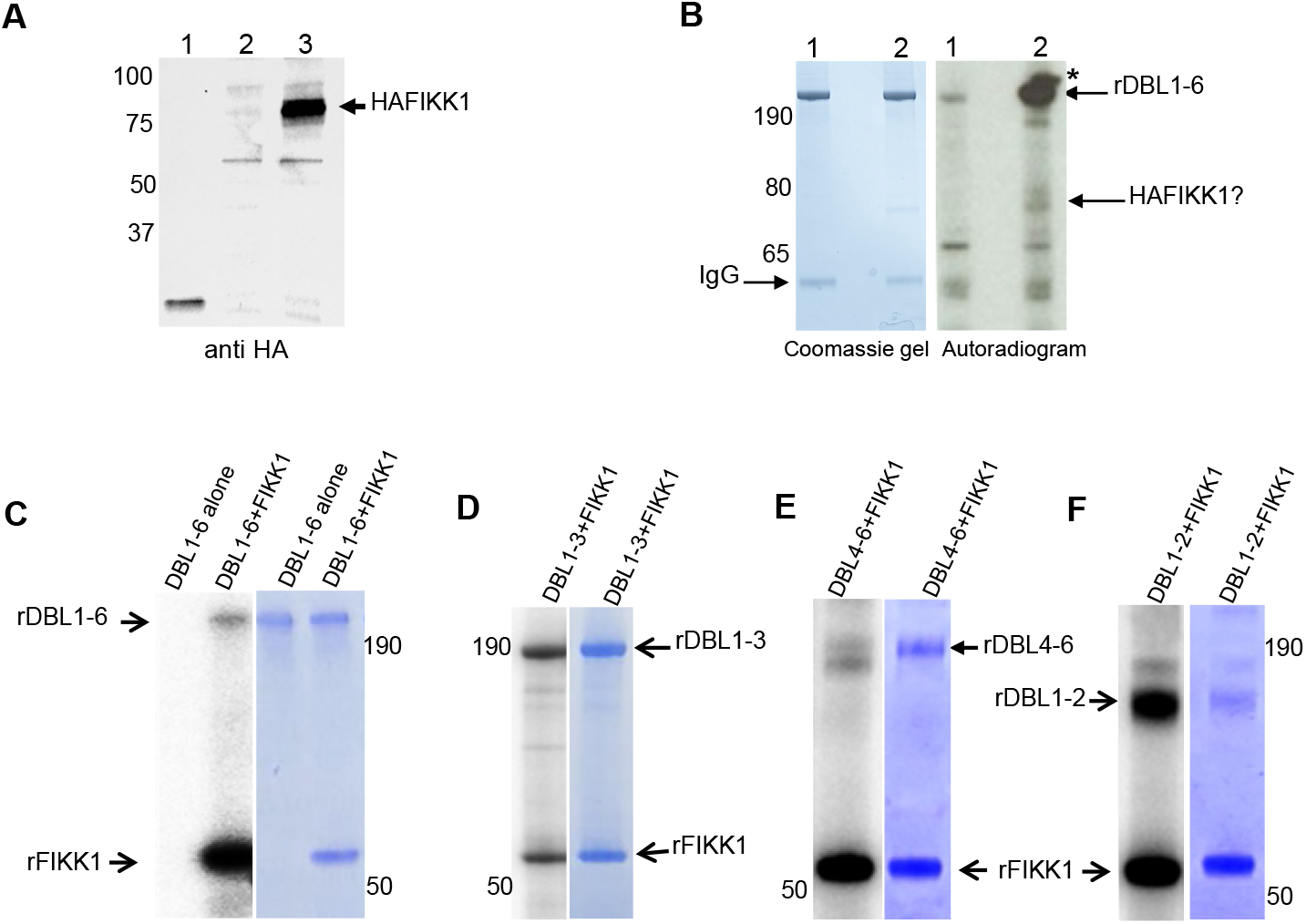
FIKK1 phosphorylates VAR2CSA. **A**. Western-blot analysis using a rabbit anti-HA antibody; Immunoprecipitation experiments with DMSO- and RAPA-treated IEs total protein lysates were performed using a mouse anti-HA antibody. Lane 1: positive HA control. Lane 2: immunoprecipitation from RAPA-treated lysates; lane 3: immunoprecipitation from DMSO-treated lysates control. **B**. *In vitro* phosphorylation of rDBL1-6 by FIKK1::HA. Immunoprecipitated material from both lysates were used in a standard [γ - 32P] ATP phosphorylation kinase assay with 1µg of rDBL1-6. The reactions were loaded on a gel and stained with Coomassie (left panel) prior being exposed for autoradiography (right panel). Coomassie-stained gel is shown to ensure that equivalent amount of the exogenous substrate is present in both phosphorylation assays (Fig 5B, left panel). rDBL1-6 is indicated with an arrow. Lane 1: immunoprecipitation from RAPA-treated lysates; lane 2: immunoprecipitation from DMSO-treated control lysates. **C**. rDBL1-6 full length was used alone without kinase and as a substrate for the recombinant kinase domain of FIKK1 in the presence of [γ^32^P] ATP. **D**. Kinase assay carried out with rFIKK1KD and VAR2CSA multidomain DBL1-3. **E**. Kinase assay carried out with multidomain DBL4-6. **F**. Kinase assay carried out with multidomain DBL1-2. Autoradiograms are shown on the left panels and the corresponding Coomassie-stained gels are on the right panels. Proteins are indicated in the figure with arrows.

The above results strongly suggest that endogenous FIKK1 can phosphorylate the extracellular region of recombinant VAR2CSA. To ascertain this, and to rule out indirect phosphorylation, *in vitro* radiolabeled kinase reactions were carried out with recombinant FIKK1 kinase domain and rDBL1-6 as a substrate. We indeed confirmed that VAR2CSA rDBL1-6 is phosphorylated by the recombinant enzyme (**Fig 5C, left panel)**. No phosphorylation of VAR2CSA was observed in the negative control without FIKK1. To determine which domains of VAR2CSA are targeted, phosphorylation assays were carried out on multi-domains rDBL1-3, and rDBL4-6 corresponding respectively to the N-terminal and C-terminal multi-domains of the VAR2CSA extracellular region. Autoradiograms reveal that the radiolabeled band corresponding to the rDBL1-3 was much more intense than the signal for rDBL4-6 **(Fig 5D and E, left panels)**. Similar amount of proteins was used in the assays as verified by Coomassie-stained gel prior exposure (**Fig 5D and E, right panels)**. We further narrowed down the targeted region using the multi-domains DBL1X-2X. Autoradiogram reveal that DBL1-2 displays a strong phosphorylation signal **(Fig 5F)**. These results suggest that FIKK1-mediated VAR2CSA phosphorylation occurs mainly in the N terminal domain of the adhesin. To investigate whether FIKK1 activity could enhance VAR2CSA adhesion properties, we assessed the binding of rDBLVAR2CSA to CSA in presence or not of FIKK1 kinase plus ATP. Interestingly in 3 independent experiments, we observed a significant increase in the binding to CSA when rVAR2CSA at 3 different concentrations was preincubated with the kinase plus ATP compared to the condition without kinase plus ATP, indicating that FIKK1 mediated VAR2CSA phosphorylation enhances binding to CSA *in vitro* **(Fig 6A)**

**Fig 6.**
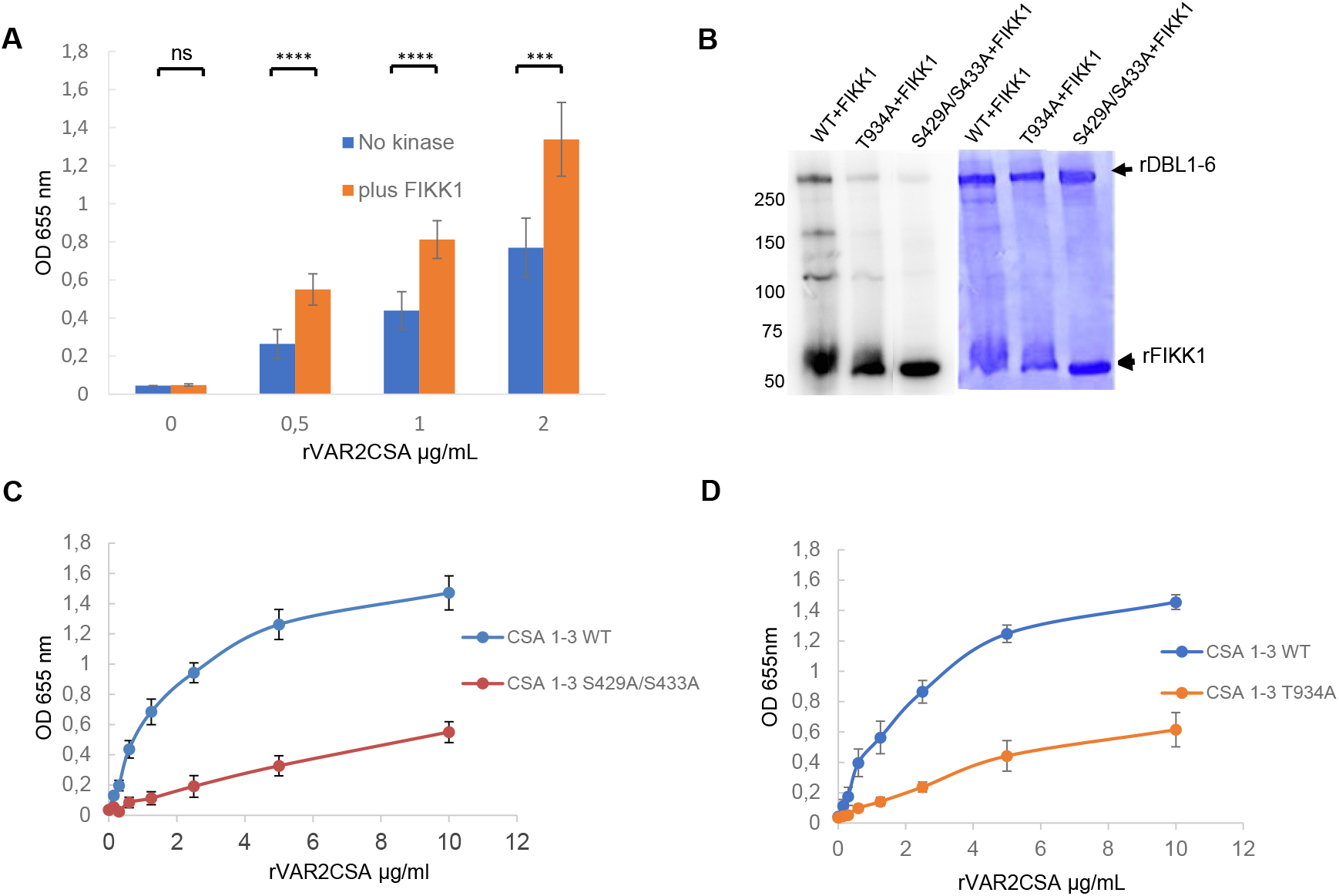
Phosphorylation increases the binding of VAR2CSA to CSA. Identification and validation of targeted VAR2CSA domains and phosphosites. **A**. FIKK1 kinase plus ATP increases binding of rVAR2CSA to CSA. 0, 0.5, 1 and 2 μg/mL of VAR2CSA recombinant proteins were preincubated 30 min at 30°C in kinase buffer supplemented with 10μMATP with 400 ng or without FIKK1 kinase and added to wells previously coated with CSA. Error bars correspond to SD between 3 independent experiments. Experimental data were analyzed using GraphPad Prism software. Statistics (paired t test; ****p = 0.0001; for 0.5, 1µg/ml and ***p= 0.0005 for 2μg/ml rDBL1-2VAR2CSA; CSA chondroitin sulfate A; OD optical density at 655nm; SD standard deviation. **B**. Effect of mutations T934A and S429A/S433A on rDBL1-6 phosphorylation by rFIKK1 kinase domain. The phosphorylation of the wild-type and mutated rDBL1-6, DBL1-6 T934A and DBL1-6 S429A/S433A proteins were tested in a standard *in vitro* kinase assay in the presence of [γ-32P] ATP with the recombinant FIKK1 kinase domain. **C**. Binding assay of wild-type DBL1-3 and mutated DBL1-3 S429A/S433A recombinant proteins to CSA-coated plates assayed by ELISA. **D**. Wild-type DBL1-3 and mutated DBL1-3 T934A recombinant proteins were assayed by ELISA for *in vitro* binding to CSA coated plates. Increasing concentrations of recombinant DBL1-3 proteins at serial dilutions of 0.156 to 10 μg/mL were added to wells previously coated with CSA. Error bars correspond to SD between 3 independent experiments. Each experiment was performed in triplicate.

### Identification and validation of targeted VAR2CSA phosphosites

To identify the phosphosites targeted by the FIKK1 kinase, a liquid chromatography-tandem mass spectrometry (LC-MS/MS) analysis was performed after an *in vitro* phosphorylation assay with rFIKK1, cold ATP and with rVAR2CSA DBL1-3 alone as a control. The analysis revealed the presence of a phosphoserine at position 433 in rDBL1-3 alone and the presence of several phosphoamino acids such as T934 and S429 after incubation with FIKK1KD **(Table1)**. These two phosphosites were reported previously to be phosphorylated in native VAR2CSA proteins [10]. Threonine 1006 and Threonine 1007 and, interestingly, Tyrosine residues are phosphorylated (Y271 and Y998) *in vitro* by FIKK1. A schematic mapping of rDBL1-3 identified phosphorylation sites by rFIKK1 kinase domain is displayed in **S5 Fig**. Site-directed mutagenesis was used to validate the MS data. Radiolabeled phosphorylation assays of recombinant DBL 1-6 wild type, S429A/S433A double mutant and T934A mutant by rFIKK1, were carried out. Interestingly, we observed a radiolabeling decrease of the VAR2CSA DBL1-6 mutant proteins compared to the DBL1-6 wild-type substrate **(Fig 6B left panel)** suggesting that T934 and S429 could be dominant phosphorylated residues by the FIKK1. We ensured that the same amounts of kinase and rVAR2CSA substrates were loaded in all reactions by a Coomassie staining of the gel prior autoradiography **(Fig 6B right panel)**. DBL1-3 WT, DBL1-3 T934A and DBL1-3 S429A/S433A recombinant proteins were expressed in HEK293 cells and purified as described previously [10]. To confirm the importance of these phosphosites in CSA adhesion, ELISA-based binding assays to CSA were carried out with increasing concentrations of wild-type and mutant rVAR2CSA. Proteins concentrations were normalized prior CSA-binding assays. A significant reduction in CSA binding was observed in 3 Independent experiments, with both mutant proteins in comparison to the wild-type protein **(Fig 6C and D**), confirming the importance of these residues in CSA adhesion

**Table 1.**
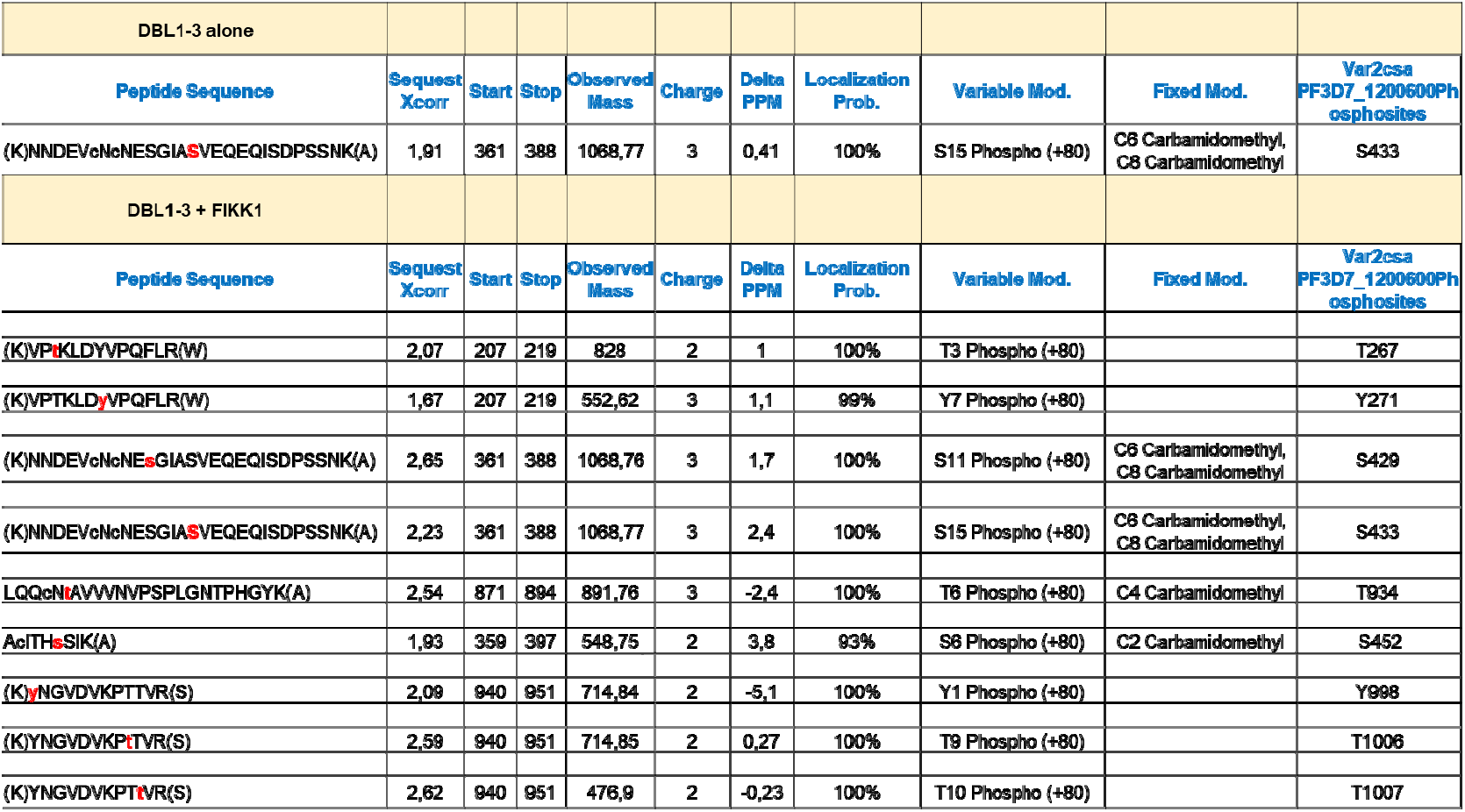
Summary of the identified rDBL1-3 phosphorylation sites after kinase assay with rFIKK1 KD.

## Discussion

Pathogenicity of placental malaria is caused by the ability of *P. falciparum*-IEs to adhere to the glycosaminoglycan CSA expressed on the surface of placental syncytiotrophoblasts. The VAR2CSA adhesin mediates the binding to CSA and stands today as the leading candidate for a vaccine aimed at protecting pregnant women malaria endemic areas against the severe clinical outcomes of placental malaria [29] [30].

It is well established that the phosphorylation of cell surface proteins is an important, reversible post-translational modification that can regulate cellular ligand interactions. We have provided in our previous study [10] direct evidence that the extracellular domain of VAR2CSA is phosphorylated (including in the CSA binding region), and that phosphatase treatment of intact IEs decreases their adhesive properties to placental CSA. This suggests that kinases play a crucial role in placental malaria virulence, and is consistent with our recent observation that both erythrocyte and *P. falciparum* casein kinase 2 enzymes phosphorylate the extracellular region of VAR2CSA and play a role in trafficking and cytoadhesion to CSA [28].

Therefore, one important task is to identify specific *Plasmodium falciparum* kinases involved in parasite multiplication or IEs cytoadhesion to host cells. Host RBC remodeling is in part due to the activity of *Plasmodium*-encoded proteins such as the PHIST proteins [31] and rifins [32] that are secreted into the RBC cytoplasm in distinct locations. In addition, several members of the intriguing FIKK kinase family are known to be exported into RBC and to be linked to pathogenicity [24].

Here, we report that following rapamycin treatment, FIKK1 knocked-out clone F5 parasites expressing VAR2CSA are significantly less adhesive to CSA than parasites carrying an intact *fikk1* locus. Since it has been reported that rapamycin could affect wild-type parasites [27], we ensured that the drug has no effect on the parental line enabling us to assign any RAPA effect to FIKK1 absence.

The impaired cytoadhesion phenotype was observed in KO FIKK1 IEs expressing similar levels of VAR2CSA than the DMSO-treated cell line, suggesting a role for this kinase in IEs cytoadhesion, possibly via the phosphorylation of VAR2CSA. Direct evidence to support phosphorylation of endogenous VAR2CSA by FIKK1 in live parasites is still missing. However, in line with this hypothesis, we showed that immunoprecipitated FIKK1-HA is able to co-immunoprecipitate with endogenous VAR2CSA and phosphorylate the recombinant extracellular VAR2CSA DBL1-6 region. Additionally, we observed a high molecular weight radiolabeled band migrating just above the radiolabeled ectodomain DBL1-6 that could correspond to the endogenous full length VAR2CSA co-immunoprecipitated with FIKK1-HA.

Interestingly, co-staining of both proteins in Maurer’s Clefts structures by IFA indicates a potential compartment where the proteins could interact. Phosphorylation of VAR2CSA by FIKK1 could occur in MCs during VAR2CSA trafficking prior final translocation into the IEs membrane. Our FIKK1 localization data consistent with recently reported results [25] indicating RBC membrane localization for FIKK1 (in addition to the Maurer’s Cleft subcellular localization). Immunoprecipitation studies indicate that native VAR2CSA was co-immunoprecipitated only from membrane fractions. Of interest, it has been shown that FIKK12 dissociates from the MC to relocate at the erythrocyte membrane at trophozoite stage and might be involved in the phosphorylation of proteins of the erythrocyte membrane skeleton [33]. It is therefore possible that a pool of FIKK1 is transferred in vesicles from the Maurer’s Cleft, trafficked along actin filaments as previously hypothesized for PfEMP1 [34, 35] and could be relocated transiently within the skeleton network and membrane periphery. Alternatively, phosphorylation could occur prior to export in the PV lumen. Either option would be compatible with FIKK1 and PfEMP1 timing of expression during the *P. falciparum* life cycle and therefore the capacity of FIKK1 to bind and phosphorylate VAR2CSA. However, elucidating the precise location and dynamics of FIKK1 and VAR2CSA interactions will require further experimental work.

VAR2CSA interaction and phosphorylation by FIKK1 was confirmed in *in vitro* experiments. Indeed, using a combination of several *in vitro* approaches (pull down, ELISA, phosphorylation assays and phosphoproteomics studies), we have demonstrated that the recombinant FIKK1 kinase domain interacts and phosphorylates rDBL1-6, mainly in the DBL1-3 region that encompass the CSA-binding domain [36, 37]. Importantly, we observed in three independent experiments a significant increase in the binding to CSA when rVAR2CSA was preincubated with the kinase plus ATP compared to the condition without kinase, indicating that FIKK1-mediated VAR2CSA phosphorylation enhances binding to CSA. These data are in line with the impaired cytoadhesion phenotype observed in KO FIKK1 IEs.

We also showed that Threonine 934, located in ID2 or CIDR domain and phosphorylated in native VAR2CSA, is targeted *in vitro* by FIKK1. We were able to validate this result by showing *in vitro* a reduced FIKK1-mediated phosphorylation of rVAR2CSA T934A mutant protein compared to the wild type substrate. Unfortunately, we were unable to show that endogenous FIKK1 targets directly this amino-acid, as we failed to raise specific and sensitive antibodies against pT934. However, in a previous study, [10] cytoadhesion of a VAR2CSA T934A transgenic IEs showed a reduced adhesion to CSA while, a phosphomimetic substitution T934D slightly enhances the binding of VAR2CSA ectodomain *in vitro* [10]. These results strengthen the argument for a critical role of this phosphosite in adhesive properties of VAR2CSA. Additionally, we showed here that Alanine substitution of Threonine 934 significantly reduces binding of DBL1-3 to CSA *in vitro*. Taken together, our present and previous data support the idea that FIKK1-mediated T934 phosphorylation regulates adhesion of IEs to placenta. Additionally, S429, a residue located in the ID1 and shown to be important for CSA binding [10], is also phosphorylated by rFIKK1.

Using Oriented Peptides Array Libraries and phosphoproteome targets, Belda *et al* have determined FIKK1 substrates motifs around T and S residues with a preference for hydrophobic aminoacids in position P+2, a feature they did not observe in other FIKKs. Interestingly, VAR2CSA contains a Valine and an Isoleucine at position P+2 respectively for T934 and S429. While the *P. falciparum* kinome is devoid of true Tyrosine kinases [14, 38], we report here that Tyrosine residues are phosphorylated by FIKK1 *in vitro*. Several FIKKs from *P. falciparum* (FIKK3, FIKK4.2, FIKK9.2 to FIKK9.5, FIKK11), as well as other kinases [39] have been shown to exhibit a dual (T/S and Y) specificity [25]. Interestingly, these include PfCK2 [40, 41], which we have shown previously to be able to phosphorylate VAR2CSA [28].

Because of their divergence from typical eukaryotic Ser/Thr kinases, in addition to their restrictive existence in Apicomplexa phylum, developing inhibitors that would selectively target the FIKKs, while not interfering with host kinases, is of high potential with respect to new therapeutic strategies. Several *in silico* approaches based on computational screenings of phytochemical and Molecular Dynamics simulations validated Rufigallol as the most potent inhibitory compound against FIKK9.5. Belda *et al* have demonstrated that all expressed members of the FIKK family can be chemically inhibited *in vitro* using a single compound, lending hope that a pan-FIKK inhibitor may enter the drug development pipeline [25].

In summary, our findings demonstrate that FIKK1 not only phosphorylates VAR2CSA but also plays a critical role in regulating the cytoadhesion of IEs to chondroitin sulfate A. Additionally, FIKK1 has been implicated in the phosphorylation of red blood cell adducin [24], a process that may further influence the adhesion phenotype observed. Together, these results advance our understanding of the molecular mechanisms governing the binding of VAR2CSA-expressing IEs to placental CSA, thereby shedding light on the processes underlying parasite sequestration in the placenta.

## Material and methods

### *P. falciparum in vitro* culture

*P. falciparum* NF54 wild-type parental line and transgenic cell lines were maintained in culture under standard conditions in O+ Human erythrocytes in RPMI 1640 containing L-glutamine (Invitrogen) supplemented with 10% Albumax I, 1 × hypoxanthine and 20 μg/mL gentamicin. IES cultures (3–5% parasitemia) at mid/late trophozoite stages were purified using CS columns on the VarioMACS system (Miltenyi). Genomic DNA extracted according to the protocol described in Blood easy extraction kit (Qiagen). Parasite cultures were regularly tested for Mycoplasma contamination (LookOut Mycoplasma PCR detection kit (Sigma)) using MSP primers. Transgenic FIKK1cKO cell line was generated from the NF54::Dicre inducible cell line to conditionally delete the kinase of interest according to the strategy previously described in [24]. The transgenic parasite population was cloned by limiting dilution.

### Conditional knockout (KO) induction of FIKK1 kinase

To induce DiCre-driven LoxP site recombination, synchronized ring-stage parasites were treated with 100 nM Rapamycin (Sigma) or dimethyl sulfoxide (DMSO) for 4 hours. Parasites were subsequently washed twice with RPMI 1640 medium and returned to culture. 48 hours later rings were harvested for qRTPCR analysis after Trizol RNA extraction. 72 hours later, mature parasites are harvested for gDNA extraction, immunofluorescence assays, MACS purified for flow cytometry and cytoadhesion assays.

### IEs cytoadhesion assays

#### Static adhesion

##### CSA coated on petri dish

20μL of CSA (1mg/mL) in PBS or PBS 1% BSA (negative control) were spotted (approximately 0.5 cm diameter circles) on a Petri dish then incubated overnight at 4°C in a humidified chamber. The spots were washed twice in PBS and blocked with PBS / BSA 1% for 1 h at RT. DMSO and RAPA-treated mature stages were purified using VarioMACS and 10^5^ IEs were added to the spotted plates at RT for 1 h. Unbound IEs were gently washed away twice with PBS. Adherent infected red blood cells were fixed with 2% glutaraldehyde and counted on 5 fields in duplicate spots with a Nikon Eclipse Ti microscope with a 10 X objective. Results are expressed as a percentage of RAPA-treated parasites binding compared to DMSO-treated parasites (100% binding). The results of static binding assay were statistically tested by Graph Prism (n=3 biological replicates)

##### 96-wells plate CSA binding assay

Inhibition of IEs binding to CSA was assessed in a 96-well plate as described in [42]. Briefly, 100µL/well of a solution of CSA (1mg/mL in PBS) or BSA (1% in PBS) was coated on a MaxiSorp™ high protein-binding capacity polystyrene 96-well plate overnight at 4°C. Plates were then washed three times with 200 μL RPMI 1640 supplemented with 2% Fetal Bovine Serum (FBS). DMSO and RAPA-treated mature sages were MACS-purified and then added into the pre-coated wells (10^6^ cells in 100 μL/well) and incubated for 1 h at 37 °C. Plates were then washed three times with 200 μL PBS. Infected erythrocytes remaining attached to the surface were lysed by addition of 50 μL 3,3⍰,5,5⍰-Tetramethylbenzidine (TMB) (Biorad) followed by 30 s vortexing on a MixMate® mixer (1000 rpm). Monitoring of the pseudo peroxidase activity of the remaining hemoglobin contained in infected erythrocytes was performed by reading the absorbance at 655 nm using a TECAN absorbance reader (Biorad). Results were expressed as a percentage of RAPA-treated binding compared to the DMSO control (100% binding). The results of 96-wells static binding assay were statistically tested by Graph Prism (n=3 biological replicates; the average binding for each experiment corresponds to 4 OD655nm readings).

##### Flow adhesion assay

For measuring flow adhesion to CSA, DMSO and RAPA FIKK1 treated parasites were used with the method previously described [26]. Briefly, IbiTreat µ-slide VI0.4 slides (ibidi) were coated with 10 mg/mL CSA for 2 hr then washed with 50 mL PBS. DMSO or RAPA-treated FIKK1 iRBC were percoll-enriched, divided in two and stained with either Hoechst 33342 (New England Biolabs) or SYBRGreen, incubated for 10 minutes, then washed in complete media. Hoechst-stained RAPA-treated parasites were mixed in a 1:1 ratio with SYBRGreen-stained DMSO-treated parasites, and vice versa. The mixed samples were flowed over the CSA-coated slides at a flow rate of 4.5 mL/hr (tumbling speed) for 10 minutes, before gradually increasing the flow rate to 150 mL/hr over 10 minutes. At each flow rate, 15 frames were imaged across the channel of the slide with a Nikon Microscope. The number of Hoechst or SYBRgreen-strained parasites in each frame was counted automatically using ImageJ, first filtering for cells then quantifying the blue/green fluorescence within each cell area. A ‘before’ slide was also generated by applying the mixed parasites to the slides with no flow. The ratio between RAPA /DMSO-treated parasites was calculated at each timepoint and used to calculate the change in adhesion frequency upon FIKK1 KO.

##### Recombinant proteins

Expression and purification of multi-domains and full-length VAR2CSA recombinant proteins DBL1-3, DBL4-6 and DBL1-6 VAR2CSA were cloned into pTT3 vector and expressed in HEK293-F (Human Embryonic Kidney) cells as soluble proteins secreted in the culture medium. Proteins were purified on a His-Trap Ni affinity column, followed by an ion exchange chromatography (SP Sepharose) and a gel filtration chromatography (Superdex 200) [28]. Site-directed mutagenesis to introduce the required mutations in DBL1-3 was carried out using the Quick changeII XL kit (Agilent France) according to the manufacturer’s protocol. Presence of mutations was verified by sequencing before protein expression (GATC). Oligonucleotides carrying the desired mutations were already described in our previous study [10]

Multidomain DBL1-2 was expressed in *E. Coli* as an His recombinant protein and purified as described previously [39]. Recombinant FIKK1 kinase domains were expressed as a tagged protein in *E. coli* and purified [25]. Recombinant protein His VHH was a gift from C. Hattab.

##### ELISA binding assay

The binding assay was carried out as previously described [10]. 100µL/well of CSA (1mg/mL) was coated in a 96-wells plate. Wells were blocked with 150 μL of PBS 1% BSA buffer per well for 2 h at 37°C. After removal of the blocking solution, 100 μL of increasing concentrations of wild type and mutated VAR2CSA recombinant proteins (0 to 10µg/mL diluted in BSA 1% 0.05% Tween 20 in PBS) were added to each well. After 3 washes in PBS 0.05% Tween (PBST), 100µL of anti-His HRP-conjugated antibody (1/3000, Qiagen) was added for 1 hour. ELISA in presence of FIKK1 or not plus ATP 10µM was carried out as described previously except that for detection a mouse monoclonal VAR2CSA antibody (10µg/ml) was added for 1 hour followed by a goat anti mouse HRP conjugated antibody (1/3000) After 3 washes with PBST, the reaction was revealed with 100 μL per well of TMB until saturation was reached. Binding was related to the level of absorbance monitored at 655 nm.

#### *In vitro* interaction assay

##### Pull down experiments

A mixture of 3μg of each recombinant His protein (His rVAR2CSA DBL1-6 + rFIKK1KD and His rVAR2CSA + rHis VHH) was incubated at 4°C for 30 min in binding buffer: 20mM Tris-HCL (pH 7.5), 0.2M NaCl, 0.1% Nonidet P40 (IGEPAL) and 10% glycerol according to protocol previously described [42]. Rabbit anti-VAR2CSA antibody bound to Dynabeads was added to each reaction mixture. The tubes were rotated at 4°C for 1 hour and the beads were recovered by centrifugation and washed four times in binding buffer. Laemmli buffer was added to the beads and heated at 95°C. Samples were separated by SDS-PAGE on 4-15% acrylamide stained-free gels prior western blotting.

##### ELISA

rVHH or rDBL 1-6 were diluted in PBS, coated on the plate at 2µg/mL and incubated overnight at 4°C. Wells were blocked with 150 μL of PBS 1% BSA buffer per well for 2 h at 37°C. After removal of the blocking solution, rDBL1-6 diluted at 2µg/mL in 1% BSA 0.05% Tween 20 PBS was added to the wells. After 3 washes in PBST, 100µL of a mouse monoclonal anti-VAR2CSA antibody (10µg/mL) was added for 1 hour. After 3 washes with PBST, samples were incubated with a goat anti-mouse HRP-conjugated antibody (dilution 1/3000). After 3 additional washes, the reaction was revealed with 100 μL per well of TMB until saturation was reached. Binding was related to the level of absorbance monitored at 655 nm.

##### *Var* gene expression analysis by qPCR

RNA from transgenic NF54CSA synchronized ring stages parasites was extracted with TRIzol (Qiagen) following manufacturer’s instructions. cDNA synthesis was performed using random primers after DNase I treatment (TURBO DNase, Ambion) using the SuperScript III First Stand Synthesis system (Invitrogen). Primer pairs used to detect each *var* gene expression have been described previously [14]. Real-time PCR reactions were performed on a CFX 96 thermocycler (Biorad). Transcriptional level of each *var* gene was normalized with the housekeeping control gene seryl tRNA transferase (PlasmoDB: PF3D7_0717700).

##### VAR2CSA surface expression analysis by flow cytometry analysis

VarioMACS (Miltenyi Biotec France) purified IEs at mid and late trophozoite stages were resuspended in PBS 0.2% BSA and counted. For each assay, 0.5 × 10^6^ IEs were washed in PBS and incubated with 50 μL of purified rabbit polyclonal anti-VAR2CSA IgG diluted 1/100 in PBS 0.2% BSA for 1h at RT. IEs were washed twice with PBS 0.2% BSA and resuspended in 100μL of PE-conjugated donkey anti-rabbit IgG diluted 1/100 in PBS 0.2% BSA for 30 min at RT. After two washes in PBS 0.2% BSA, IEs were fixed in 4% paraformaldehyde in PBS and kept at 4°C overnight in the dark. Cells were then washed twice with PBS and parasite nuclei were stained with TO-PRO-3 dye (1/10,000 dilution). Samples were analyzed by flow cytometry using a BD FACScanto II flow cytometer (Becton Dickinson France) and data were processed using FlowJo 10.0 software. The results are presented as the geometric mean of fluorescence intensities.

##### Preparation of total, soluble and membrane IEs protein extracts

Total protein extracts were prepared from asexual mature stages parasites harvested by MACS purification. After washing with cold PBS, the parasites pellets were resuspended in cold lysis buffer (150mM NaCl, 50mM Tris HCl pH 8.0, 1% NP40, 0.5% Sodium Deoxycholate, 0.05% SDS supplemented with proteases and phosphatases inhibitors + 10µM ATP) incubated on ice and sonicated briefly. The lysates were cleared by centrifugation (20,000 x g for 20 min at 4°C). The total amount of proteins in the supernatant was measured using the Bradford assay, was visualized on a stain-free (from Biorad) or Coomassie-stained SDS gel or was collected for immunoprecipitation, western blot or kinase assays applications.

IEs soluble (Triton soluble) and membrane (Triton insoluble) fractions were prepared as described previously [43]. Briefly, the IEs were resuspended in NETT lysis buffer (150mM NaCl, 5mM EDTA, 50mM Tris HCl pH 8.0, 1% Triton X-100) supplemented with protease, phosphatase inhibitors and ATP. The membrane fraction was separated from soluble cytosolic material by centrifugation at 20,000 x g at 4°C for 30 min. The pellet was resuspended and dissolved at room temperature in Tris-saline buffer (pH8) supplemented with 2% SDS, proteases and phosphatases inhibitors. After centrifugation at RT at 20,000 x g, the resulting supernatant corresponding to the solubilized membrane fraction was recovered.

##### Western blot

Anti-HA western blots were performed using a mouse monoclonal immuno-purified antibody following manufacturer’s recommendations (Biolegend; 1/1000 dilution) followed by a secondary goat anti-mouse antibody conjugated to peroxidase (1/4000). A rabbit anti-HA from Clontech (1/1000) was used as well followed by a secondary goat anti-rabbit HRP conjugated antibody (1/4000). For VAR2CSA detection, western blots were carried out using a goat polyclonal anti-VAR2CSA antibody (1/1000) followed by an incubation with a secondary rabbit anti-goat antibody conjugated to peroxidase (1/4000).

##### Immunoprecipitation

Soluble or membrane fractions lysates of IEs were incubated for 2 h with 3μg of mouse IgG isotype or mouse immuno-purified anti-HA (Biolegend clone 16B12) (Santa Cruz Biotechnologies). Immuno-conjugated material was recovered with 20μL of protein G Sepharose beads after centrifugation, washed four times in NETT buffer then in Tris buffer saline and samples were heated for 3 min at 95°C for western blot applications. For kinase assays, the beads were washed once with kinase assay buffer and used as a source of kinase in standard phosphorylation assays.

##### Kinase assays

Immunoprecipitated material bound to agarose beads or purified recombinant kinase domain of FIKK1[25] were used as a source of kinase in a standard reaction (30 µL) using 25 mM Tris/HCl, pH 7.5, 20 mM MgCl2, 2mM MnCl2, 10µM ATP in the presence of 2.5 µCi ATP, [γ−32P]. A mix of kinase buffer plus rFIKK1 was performed prior adding cold and hot ATP and rVAR2CSA substrates. The reactions were proceeded for 30 min at 30 °C and were stopped by the addition of Laemmli buffer, boiled for 3 min and analyzed by SDS PAGE. The gels were dried and submitted to autoradiography. Cold kinase assay with recombinant proteins was performed as above with cold ATP only.

##### Immunofluorescence assays

HA-FIKK1 protein expression was studied by immunofluorescence assays on cold-methanol/ acetone-fixed infected erythrocytes culture smears with a mouse anti-HA followed by an anti-mouse secondary antibody conjugated to Alexa Fluor 568. Double labelling experiments were performed as follows: mouse anti-HA antibody (1/200 dilution) and rat anti-PfSBP1 were incubated with PBS, 1% BSA, for 1 h. After three washes in PBS, an Alexa Fluor 568 conjugated donkey anti-mouse antibody (1/400 dilution) and an Alexa Fluor 488 donkey ant-rat antibody (1/400 dilution) were incubated for 1 hour.

VAR2CSA surface localization was analyzed on live cells with a specific rabbit anti-VAR2CSA antibody (1/400) as described previously [28]. Co-staining with PfSBP1 was performed on fixed smears as above with a goat anti-VAR2CSA and a mouse anti-HA followed by an Alexa Fluor 568 anti-goat and Alexa Fluor 488 anti-rat antibodies. Co-staining of VAR2CSA and FIKK1 proteins was assessed as followed: smears were incubated with a mouse anti-HA and a goat anti-VAR2CSA diluted as above, washed three times, then incubated respectively with an Alexa Fluor 568 anti-mouse and an Alexa Fluor 488 anti-goat antibodies (1/400 dilution). The slides were washed, incubated with Hoechst 33342 dye (1/1000 in PBS) for nuclei staining for 5 min, washed and mounted with Fluoromount. Labelled specimens were examined with a Digital camera on a LSM700 Zeiss microscope.

#### Mass spectrometry

##### Sample preparation

Each sample was diluted in 50 µL of 4 M urea + 10% acetonitrile and buffered with Tris-HCl pH 8.5 at a final concentration of 30 mM. Reduction was performed with 10mM dithioerythritol at 37°C for one hour with constant shaking (600 rpm). Samples were again buffered to pH 8.5 with Tris pH 10-11 and alkylation was performed with 40 mM iodoacetamide at 37°C for 45 min with constant shaking in a light-protected environment. Reactions were quenched by the addition of dithioerythritol to a final concentration of 10 mM. Samples were then diluted five-fold with 50 mM ammonium bicarbonate and protein digestion was performed overnight at 37°C using mass spectrometry grade Trypsin Gold (1/50 enzyme-protein) and 10 mM CaCl2 or using chymotrypsin. Reactions were stopped by the addition of 2 µL of pure formic acid. Peptides were desalted on C18 StageTips Eluted peptides were dried by vacuum centrifugation prior to LC-MS/MS injections or submitted to phosphopeptide enrichment. Selective enrichment of phosphopeptides were performed on homemade titania tips. Prior to sample loading, the titania tips were equilibrated with 0.75% TFA, 60% acetonitrile, lactic acid 300mg/mL (Solution A). The digested peptides were resuspended in 20 µL of solution A and loaded on a titania tip. After a successive washing step with solution A and 0.1% TFA, 80% ACN (solution B), two elutions were performed with 0.5% ammonium hydroxide and 0.5% piperidine. Eluted fractions were acidified with FA (Formic Acid) and dried in a speedvac.

##### LC-MS/MS

Samples were resuspended in 2% acetonitrile, 0.1% FA and nano-flow separations were performed on a Dionex Ultimate 3000 RSLC nano UPLC system (Thermo Fischer Scientific) on-line connected with an Orbitrap Elite Mass Spectrometer (Thermo Fischer Scientific). A homemade capillary pre-column (Magic AQ C18; 3 µm to 200 Å; 2 cm × 100 µm ID) was used for sample trapping and cleaning. A C18 tip-capillary column (Nikkyo Technos Co; Magic AQ C18; 3 µm to 100 Å; 15 cm × 75 µm ID) was then used for analytical separations at 250 nL/min over 75 min using biphasic gradients. Samples were analysed in data-dependent acquisition mode with a dynamic exclusion of 40 sec. The twenty most intense parent ions from each MS survey scan (m/z 300-1800) were selected and fragmented by CID (Collision Induced Dissociation) into the Linear Ion Trap. Orbitrap MS survey scans resolution was set at 60,000 (at 400 m/z) and fragments were acquired at low resolution in centroid mode.

##### Data processing

Raw data were processed using SEQUEST, MS Amanda and Mascot in Proteome Discoverer v.2.4 against a concatenated database consisting of the Uniprot human reference proteome (Release 2014_06) and VAR2CSA sequence. Enzyme specificity was set to trypsin or chymotrypsin and a minimum of six amino acids was required for peptide identification. Up to two missed cleavages were allowed. For the search, carbamidomethylation was set as a fixed modification, whereas oxidation (M), acetylation (protein N-term) and Phosphorylation (S, T, Y) were considered as variable modifications. Data were further processed using X! Tandem and inspected in Scaffold 5.1 (Proteome Software, Portland, USA). Spectra of interest were manually validated. Peptide-spectrum matches with Mascot score > 18 and/or SEQUEST score > 3 were considered as correctly assigned.

## Data Availability

All data are found in the manuscript and Supporting information files

## Supporting information figure legends

Supplementary figure 1: ELISA showing that recombinant FIKK1 kinase domain (KD) interacts with recombinant DBL1-6 VAR2CSA and not with an irrelevant protein. VHH, FIKK1 or PBS were coated on the plate and then rVAR2CSA was added. Detection was performed with a monoclonal anti-VAR2CSA followed by an anti-mouse HRP-conjugated antibody. After addition of TMB, absorbance was monitored at 650 nm. Statistics (n=3; Unpaired t test *** p=0.0006).

**Supplementary figure 2: PCR genotyping of genomic DNA of NF54 WT parental line, DMSO- and RAPA-treated parasites**. Diagnostic for 5’ integration, 3’ integration and wild type locus: lane 1: NF54WT; lane 2: DMSO FIKK1::HA; lane 3: H20; lane 4: MW. Diagnostic for excision locus: lane 4: DMSO FIKK1::HA; Lane 5: RAPA FIKK1::HA; lane 6: NF54WT; lane 7: H20; Lane 8: MW.

**Supplementary figure 3: VAR2CSA IEs surface expression**. Flow cytometry histograms showing PE fluorescence intensity used to determine the percentage of IEs expressing VAR2CSA in each transgenic line. The histogram shown in the left panel is an illustration of one of the 3 independent experiments giving similar results. Representation of the geometrical mean of fluorescence intensity in RAPA-treated parasites compared to the control (DMSO-treated) from 3 independent experiments is indicated in the right panel.

**Supplementary figure 4: Rapamycin treatment does not impair VAR2CSA expression and cytoadhesion of the NF54 wild type parental line**. **A**. VAR2CSA transcripts profile of the RAPA-treated NF54WT parental line analyzed by qRTPCR. **B**. Flow cytometry histograms showing PE fluorescence intensity to determine VAR2CSA surface expression of DMSO and RAPA-treated NF54VAR2CSA parental cell line using anti-VAR2CSA antibodies. **C**. Geometric means of fluorescence intensities is indicated. **D**. Static cytoadhesion assay on Petri dish. **E**. Static adhesion assay in 96 wells plate.

Supplementary figure 5: Schematic view of rDBL1-3 identified sites phosphorylated by the rFIKK1 kinase domain

## References

1. World malaria report 2025 2025. Available from: https://www.who.int/publications/i/item/9789240117822

2. Feachem RGA, Chen I, Akbari O, Bertozzi-Villa A, Bhatt S, Binka F, et al. Malaria eradication within a generation: ambitious, achievable, and necessary. Lancet. 2019;394(10203):1056–112. Epub 2019/09/13. doi: 10.1016/S0140-6736(19)31139-0. PubMed PMID: 31511196.

3. Abu Bonsra E, Amankwah Osei P, Adjei Kyeremeh E, Adama S, Sekyi AG, King EF. Factors associated with malaria in pregnancy among women attending ANC clinics in selected districts of the Ashanti Region, Ghana. Malar J. 2025;24(1):8. Epub 2025/01/12. doi: 10.1186/s12936-025-05244-6. PubMed PMID: 39799328; PubMed Central PMCID: PMCPMC11724469.

4. Storm J, Jespersen JS, Seydel KB, Szestak T, Mbewe M, Chisala NV, et al. Cerebral malaria is associated with differential cytoadherence to brain endothelial cells. EMBO Mol Med. 2019;11(2). Epub 2019/01/06. doi: 10.15252/emmm.201809164. PubMed PMID: 30610112; PubMed Central PMCID: PMCPMC6365927.

5. Voigt S, Hanspal M, LeRoy PJ, Zhao PS, Oh SS, Chishti AH, et al. The cytoadherence ligand Plasmodium falciparum erythrocyte membrane protein 1 (PfEMP1) binds to the P. falciparum knob-associated histidine-rich protein (KAHRP) by electrostatic interactions. Mol Biochem Parasitol. 2000;110(2):423–8. Epub 2000/11/09. doi: 10.1016/s0166-6851(00)00281-4. PubMed PMID: 11071296.

6. Fried M, Duffy PE. Adherence of Plasmodium falciparum to chondroitin sulfate A in the human placenta. Science. 1996;272(5267):1502–4. Epub 1996/06/07. doi: 10.1126/science.272.5267.1502. PubMed PMID: 8633247.

7. Fried M, Domingo GJ, Gowda CD, Mutabingwa TK, Duffy PE. Plasmodium falciparum: chondroitin sulfate A is the major receptor for adhesion of parasitized erythrocytes in the placenta. Exp Parasitol. 2006;113(1):36–42. Epub 2006/01/25. doi: 10.1016/j.exppara.2005.12.003. PubMed PMID: 16430888.

8. Umbers AJ, Boeuf P, Clapham C, Stanisic DI, Baiwog F, Mueller I, et al. Placental malaria-associated inflammation disturbs the insulin-like growth factor axis of fetal growth regulation. J Infect Dis. 2011;203(4):561–9. Epub 2011/01/11. doi: 10.1093/infdis/jiq080. PubMed PMID: 21216864; PubMed Central PMCID: PMCPMC3071224.

9. Klement E, Medzihradszky KF. Extracellular Protein Phosphorylation, the Neglected Side of the Modification. Mol Cell Proteomics. 2017;16(1):1–7. Epub 2016/11/12. doi: 10.1074/mcp.O116.064188. PubMed PMID: 27834735; PubMed Central PMCID: PMCPMC5217775.

10. Dorin-Semblat D, Tetard M, Claes A, Semblat JP, Dechavanne S, Fourati Z, et al. Phosphorylation of the VAR2CSA extracellular region is associated with enhanced adhesive properties to the placental receptor CSA. PLoS Biol. 2019;17(6):e3000308. Epub 2019/06/11. doi: 10.1371/journal.pbio.3000308. PubMed PMID: 31181082; PubMed Central PMCID: PMCPMC6586358.

11. Cipak L. Protein Kinases: Function, Substrates, and Implication in Diseases. Int J Mol Sci. 2022;23(7). Epub 2022/04/13. doi: 10.3390/ijms23073560. PubMed PMID: 35408921; PubMed Central PMCID: PMCPMC8998185.

12. Ahsan R, Khan MM, Mishra A, Noor G, Ahmad U. Protein Kinases and their Inhibitors Implications in Modulating Disease Progression. Protein J. 2023;42(6):621–32. Epub 2023/09/28. doi: 10.1007/s10930-023-10159-9. PubMed PMID: 37768476.

13. Arendse LB, Wyllie S, Chibale K, Gilbert IH. Plasmodium Kinases as Potential Drug Targets for Malaria: Challenges and Opportunities. ACS Infect Dis. 2021;7(3):518–34. Epub 2021/02/17. doi: 10.1021/acsinfecdis.0c00724. PubMed PMID: 33590753; PubMed Central PMCID: PMCPMC7961706.

14. Ward P, Equinet L, Packer J, Doerig C. Protein kinases of the human malaria parasite Plasmodium falciparum: the kinome of a divergent eukaryote. BMC Genomics. 2004;5:79. Epub 11. 2004/10/14. doi: 10.1186/1471-2164-5-79. PubMed PMID: 15479470; PubMed Central PMCID: PMCPMC526369.

15. Prasad MR, Trivedi V. Molecular Investigation of FIKK Kinase Family to Develop PCR-Based Diagnosis of Plasmodium falciparum. Mol Biotechnol. 2024. Epub 2024/12/02. doi: 10.1007/s12033-024-01335-y. PubMed PMID: 39621256.

16. Rajendra Prasad M, Anil Kumar D, Dodwani E, Singh S, Trivedi V. Antigenic determinant analysis of FIKK Kinase family to identify novel candidates for diagnosis of Plasmodium falciparum. Microb Pathog. 2025;209:108070. Epub 2025/10/05. doi: 10.1016/j.micpath.2025.108070. PubMed PMID: 41045976.

17. Siddiqui G, Proellochs NI, Cooke BM. Identification of essential exported Plasmodium falciparum protein kinases in malaria-infected red blood cells. Br J Haematol. 2020;188(5):774–83. Epub 2019/10/28. doi: 10.1111/bjh.16219. PubMed PMID: 31650539.

18. Nunes MC, Goldring JP, Doerig C, Scherf A. A novel protein kinase family in Plasmodium falciparum is differentially transcribed and secreted to various cellular compartments of the host cell. Mol Microbiol. 2007;63(2):391–403. Epub 2006/12/22. doi: 10.1111/j.1365-2958.2006.05521.x. PubMed PMID: 17181785.

19. Mundwiler-Pachlatko E, Beck HP. Maurer’s clefts, the enigma of Plasmodium falciparum. Proc Natl Acad Sci U S A. 2013;110(50):19987–94. Epub 2013/11/29. doi: 10.1073/pnas.1309247110. PubMed PMID: 24284172; PubMed Central PMCID: PMCPMC3864307.

20. Bekic V, Kilian N. Novel secretory organelles of parasite origin - at the center of host-parasite interaction. Bioessays. 2023;45(9):e2200241. Epub 2023/07/31. doi: 10.1002/bies.202200241. PubMed PMID: 37518819.

21. Brandt GS, Bailey S. Dematin, a human erythrocyte cytoskeletal protein, is a substrate for a recombinant FIKK kinase from Plasmodium falciparum. Mol Biochem Parasitol. 2013;191(1):20–3. Epub 2013/08/27. doi: 10.1016/j.molbiopara.2013.08.003. PubMed PMID: 23973789.

22. D AK, Shrivastava D, Sahasrabuddhe AA, Habib S, Trivedi V. Plasmodium falciparum FIKK9.1 is a monomeric serine-threonine protein kinase with features to exploit as a drug target. Chem Biol Drug Des. 2021;97(4):962–77. Epub 2021/01/25. doi: 10.1111/cbdd.13821. PubMed PMID: 33486853.

23. Kats LM, Fernandez KM, Glenister FK, Herrmann S, Buckingham DW, Siddiqui G, et al. An exported kinase (FIKK4.2) that mediates virulence-associated changes in Plasmodium falciparum-infected red blood cells. Int J Parasitol. 2014;44(5):319–28. Epub 2014/02/18. doi: 10.1016/j.ijpara.2014.01.003. PubMed PMID: 24530877.

24. Davies H, Belda H, Broncel M, Ye X, Bisson C, Introini V, et al. An exported kinase family mediates species-specific erythrocyte remodelling and virulence in human malaria. Nat Microbiol. 2020;5(6):848–63. Epub 2020/04/15. doi: 10.1038/s41564-020-0702-4. PubMed PMID: 32284562; PubMed Central PMCID: PMCPMC7116245.

25. Belda H, Bradley D, Christodoulou E, Nofal SD, Broncel M, Jones D, et al. The fast-evolving FIKK kinase family of Plasmodium falciparum can be inhibited by a single compound. Nat Microbiol. 2025;10(6):1463–83. Epub 2025/05/20. doi: 10.1038/s41564-025-02017-4. PubMed PMID: 40389650; PubMed Central PMCID: PMCPMC12137140.

26. Davies H, Belda H, Broncel M, Dalimot J, Treeck M. PerTurboID, a targeted in situ method reveals the impact of kinase deletion on its local protein environment in the cytoadhesion complex of malaria-causing parasites. Elife. 2023;12. Epub 2023/09/22. doi: 10.7554/eLife.86367. PubMed PMID: 37737226; PubMed Central PMCID: PMCPMC10564455.

27. Tiburcio M, Yang ASP, Yahata K, Suarez-Cortes P, Belda H, Baumgarten S, et al. A Novel Tool for the Generation of Conditional Knockouts To Study Gene Function across the Plasmodium falciparum Life Cycle. mBio. 2019;10(5). Epub 2019/09/19. doi: 10.1128/mBio.01170-19. PubMed PMID: 31530668; PubMed Central PMCID: PMCPMC6751054.

28. Dorin-Semblat D, Semblat JP, Hamelin R, Srivastava A, Tetard M, Matesic G, et al. Casein Kinases 2-dependent phosphorylation of the placental ligand VAR2CSA regulates Plasmodium falciparum-infected erythrocytes cytoadhesion. PLoS Pathog. 2025;21(1):e1012861. Epub 2025/01/13. doi: 10.1371/journal.ppat.1012861. PubMed PMID: 39804934; PubMed Central PMCID: PMCPMC11761665.

29. Sirima SB, Richert L, Chene A, Konate AT, Campion C, Dechavanne S, et al. PRIMVAC vaccine adjuvanted with Alhydrogel or GLA-SE to prevent placental malaria: a first-in-human, randomised, double-blind, placebo-controlled study. Lancet Infect Dis. 2020;20(5):585–97. Epub 2020/02/08. doi: 10.1016/S1473-3099(19)30739-X. PubMed PMID: 32032566.

30. Tomlinson A, Semblat JP, Gamain B, Chene A. VAR2CSA-Mediated Host Defense Evasion of Plasmodium falciparum Infected Erythrocytes in Placental Malaria. Front Immunol. 2020;11:624126. Epub 2021/02/27. doi: 10.3389/fimmu.2020.624126. PubMed PMID: 33633743; PubMed Central PMCID: PMCPMC7900151.

31. Warncke JD, Vakonakis I, Beck HP. Plasmodium Helical Interspersed Subtelomeric (PHIST) Proteins, at the Center of Host Cell Remodeling. Microbiol Mol Biol Rev. 2016;80(4):905–27. Epub 2016/09/02. doi: 10.1128/MMBR.00014-16. PubMed PMID: 27582258; PubMed Central PMCID: PMCPMC5116875.

32. Wahlgren M, Goel S, Akhouri RR. Variant surface antigens of Plasmodium falciparum and their roles in severe malaria. Nat Rev Microbiol. 2017;15(8):479–91. Epub 2017/06/13. doi: 10.1038/nrmicro.2017.47. PubMed PMID: 28603279.

33. Nunes MC, Okada M, Scheidig-Benatar C, Cooke BM, Scherf A. Plasmodium falciparum FIKK kinase members target distinct components of the erythrocyte membrane. PLoS One. 2010;5(7):e11747. Epub 2010/07/30. doi: 10.1371/journal.pone.0011747. PubMed PMID: 20668526; PubMed Central PMCID: PMCPMC2909202.

34. Przyborski JM, Nyboer B, Lanzer M. Ticket to ride: export of proteins to the Plasmodium falciparum-infected erythrocyte. Mol Microbiol. 2016;101(1):1–11. Epub 2016/03/22. doi: 10.1111/mmi.13380. PubMed PMID: 26996123.

35. Behl A, Kumar V, Bisht A, Panda JJ, Hora R, Mishra PC. Cholesterol bound Plasmodium falciparum co-chaperone ‘PFA0660w’ complexes with major virulence factor ‘PfEMP1’ via chaperone ‘PfHsp70-x’. Sci Rep. 2019;9(1):2664. Epub 2019/02/26. doi: 10.1038/s41598-019-39217-y. PubMed PMID: 30804381; PubMed Central PMCID: PMCPMC6389991.

36. Dechavanne S, Srivastava A, Gangnard S, Nunes-Silva S, Dechavanne C, Fievet N, et al. Parity-dependent recognition of DBL1X-3X suggests an important role of the VAR2CSA high-affinity CSA-binding region in the development of the humoral response against placental malaria. Infect Immun. 2015;83(6):2466–74. Epub 2015/04/01. doi: 10.1128/IAI.03116-14. PubMed PMID: 25824842; PubMed Central PMCID: PMCPMC4432739.

37. Srivastava A, Gangnard S, Dechavanne S, Amirat F, Lewit Bentley A, Bentley GA, et al. Var2CSA minimal CSA binding region is located within the N-terminal region. PLoS One. 2011;6(5):e20270. Epub 2011/06/01. doi: 10.1371/journal.pone.0020270. PubMed PMID: 21625526; PubMed Central PMCID: PMCPMC3098292 ATIP-Avenir, the authors have declared that this did not alter their adherence to all the PLoS ONE policies on sharing data and materials.

38. Adderley J, Doerig C. Comparative analysis of the kinomes of Plasmodium falciparum, Plasmodium vivax and their host Homo sapiens. BMC Genomics. 2022;23(1):237. Epub 2022/03/30. doi: 10.1186/s12864-022-08457-0. PubMed PMID: 35346035; PubMed Central PMCID: PMCPMC8960227.

39. Low H, Chua CS, Sim TS. Plasmodium falciparum possesses a unique dual-specificity serine/threonine and tyrosine kinase, Pfnek3. Cell Mol Life Sci. 2012;69(9):1523–35. Epub 2011/11/26. doi: 10.1007/s00018-011-0888-y. PubMed PMID: 22116321; PubMed Central PMCID: PMCPMC11114921.

40. Graciotti M, Alam M, Solyakov L, Schmid R, Burley G, Bottrill AR, et al. Malaria protein kinase CK2 (PfCK2) shows novel mechanisms of regulation. PLoS One. 2014;9(3):e85391. Epub 2014/03/25. doi: 10.1371/journal.pone.0085391. PubMed PMID: 24658579; PubMed Central PMCID: PMCPMC3962329.

41. Ruiz-Carrillo D, Lin J, El Sahili A, Wei M, Sze SK, Cheung PCF, et al. The protein kinase CK2 catalytic domain from Plasmodium falciparum: crystal structure, tyrosine kinase activity and inhibition. Sci Rep. 2018;8(1):7365. Epub 2018/05/11. doi: 10.1038/s41598-018-25738-5. PubMed PMID: 29743645; PubMed Central PMCID: PMCPMC5943518.

42. Chene A, Gangnard S, Guadall A, Ginisty H, Leroy O, Havelange N, et al. Preclinical immunogenicity and safety of the cGMP-grade placental malaria vaccine PRIMVAC. EBioMedicine. 2019;42:145–56. Epub 2019/03/20. doi: 10.1016/j.ebiom.2019.03.010. PubMed PMID: 30885725; PubMed Central PMCID: PMCPMC6491931.

43. Gamain B, Dorin-Semblat D. Extraction and Immunoprecipitation of VAR2CSA, the PfEMP1 Associated with Placental Malaria. Methods Mol Biol. 2022;2470:257–71. Epub 2022/07/27. doi: 10.1007/978-1-0716-2189-9_19. PubMed PMID: 35881351.

