## Supplementary figures for "FIKK1, a member of the FIKK kinase family, phosphorylates VAR2CSA and regulates adhesion of *Plasmodium falciparum*-infected erythrocytes to the placental receptor CSA"

### Slide 1
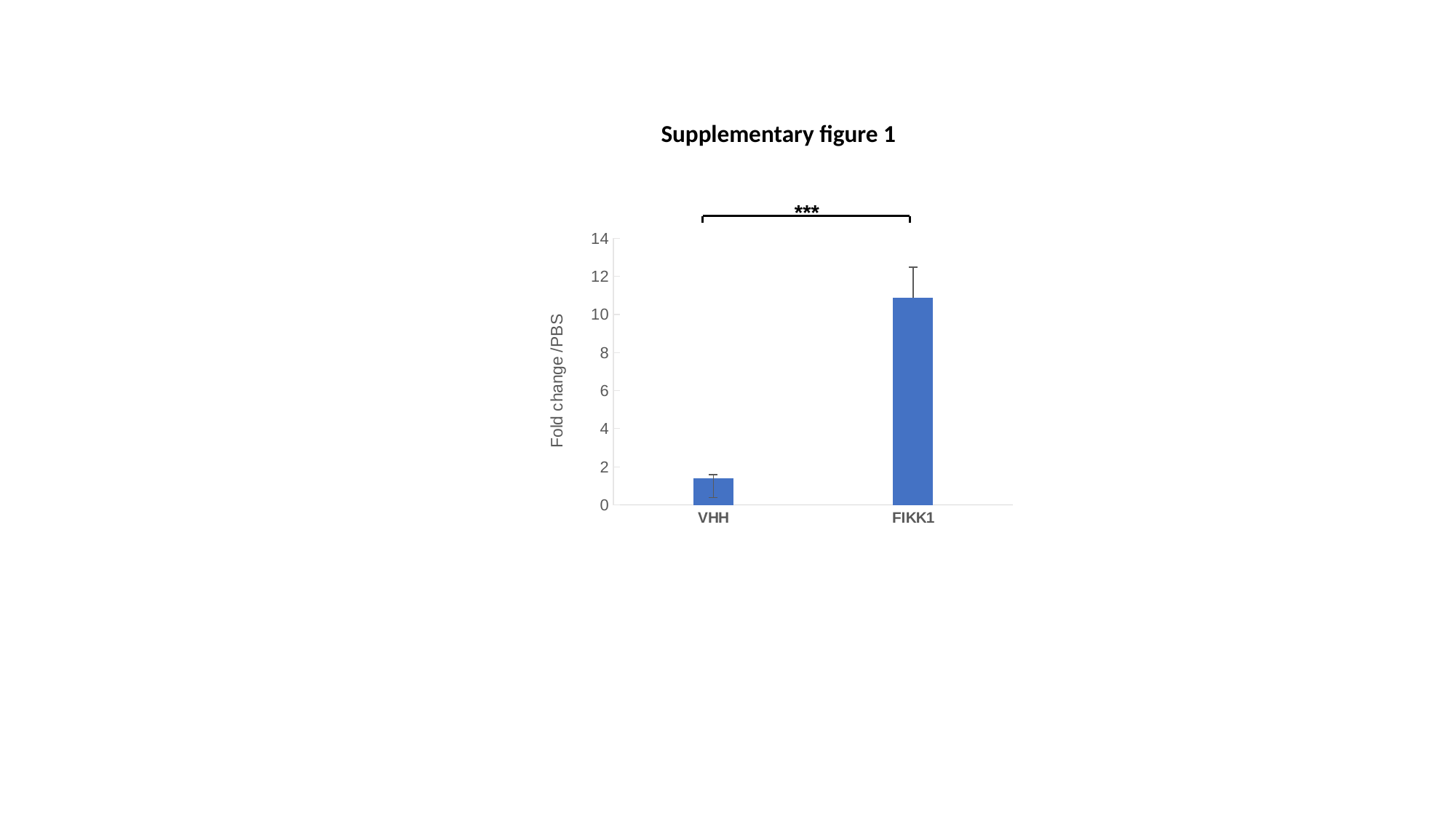

Supplementary figure 1
 ***
#### Chart
| Category | |
|---|---|
| VHH | 1.3833333333333335 |
| FIKK1 | 10.893333333333333 |

### Slide 2
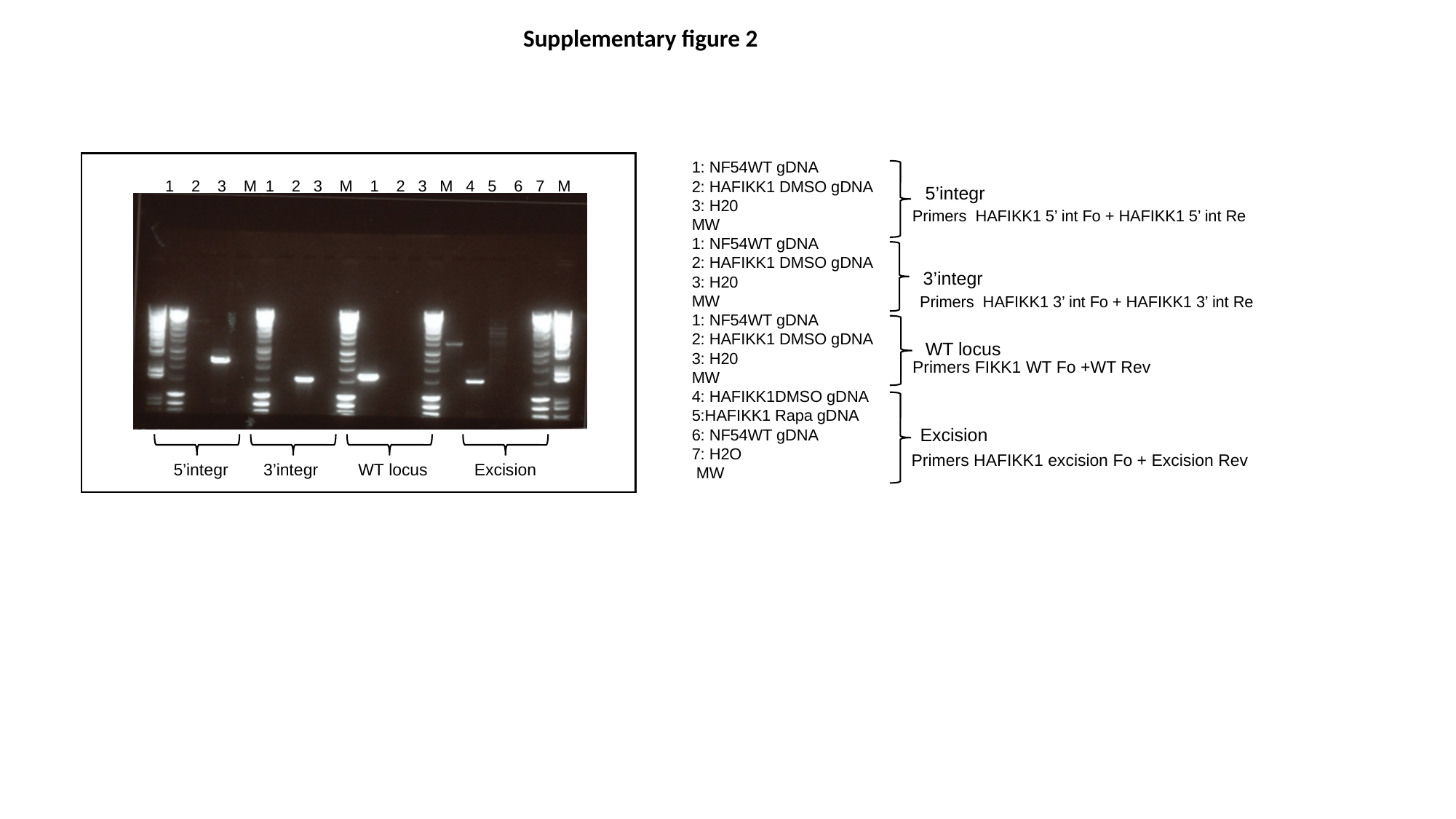

Supplementary figure 2
1: NF54WT gDNA
2: HAFIKK1 DMSO gDNA
3: H20
MW
1: NF54WT gDNA
2: HAFIKK1 DMSO gDNA
3: H20
MW
1: NF54WT gDNA
2: HAFIKK1 DMSO gDNA
3: H20
MW
4: HAFIKK1DMSO gDNA
5:HAFIKK1 Rapa gDNA
6: NF54WT gDNA
7: H2O
 MW
1 2 3 M 1 2 3 M 1 2 3 M 4 5 6 7 M
5’integr
 Primers HAFIKK1 5’ int Fo + HAFIKK1 5’ int Re
3’integr
Primers HAFIKK1 3’ int Fo + HAFIKK1 3’ int Re
WT locus
Primers FIKK1 WT Fo +WT Rev
Excision
Primers HAFIKK1 excision Fo + Excision Rev
5’integr
3’integr
WT locus
Excision

### Slide 3
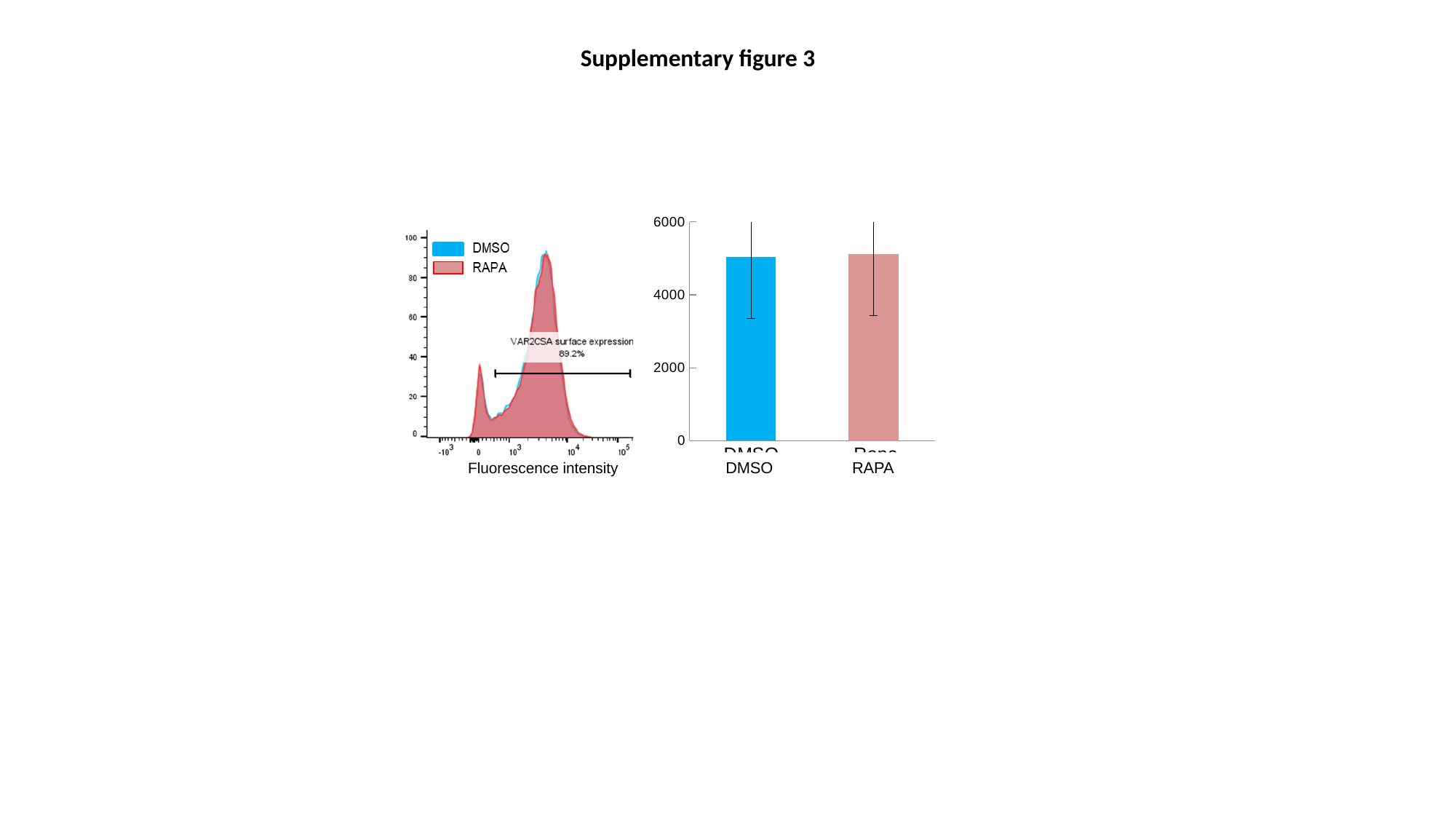

Supplementary figure 3
#### Chart
| Category | |
|---|---|
| DMSO | 5034.333333333333 |
| Rapa | 5118.0 |RAPA
DMSO
Cell count
Fluorescence intensity

### Slide 4
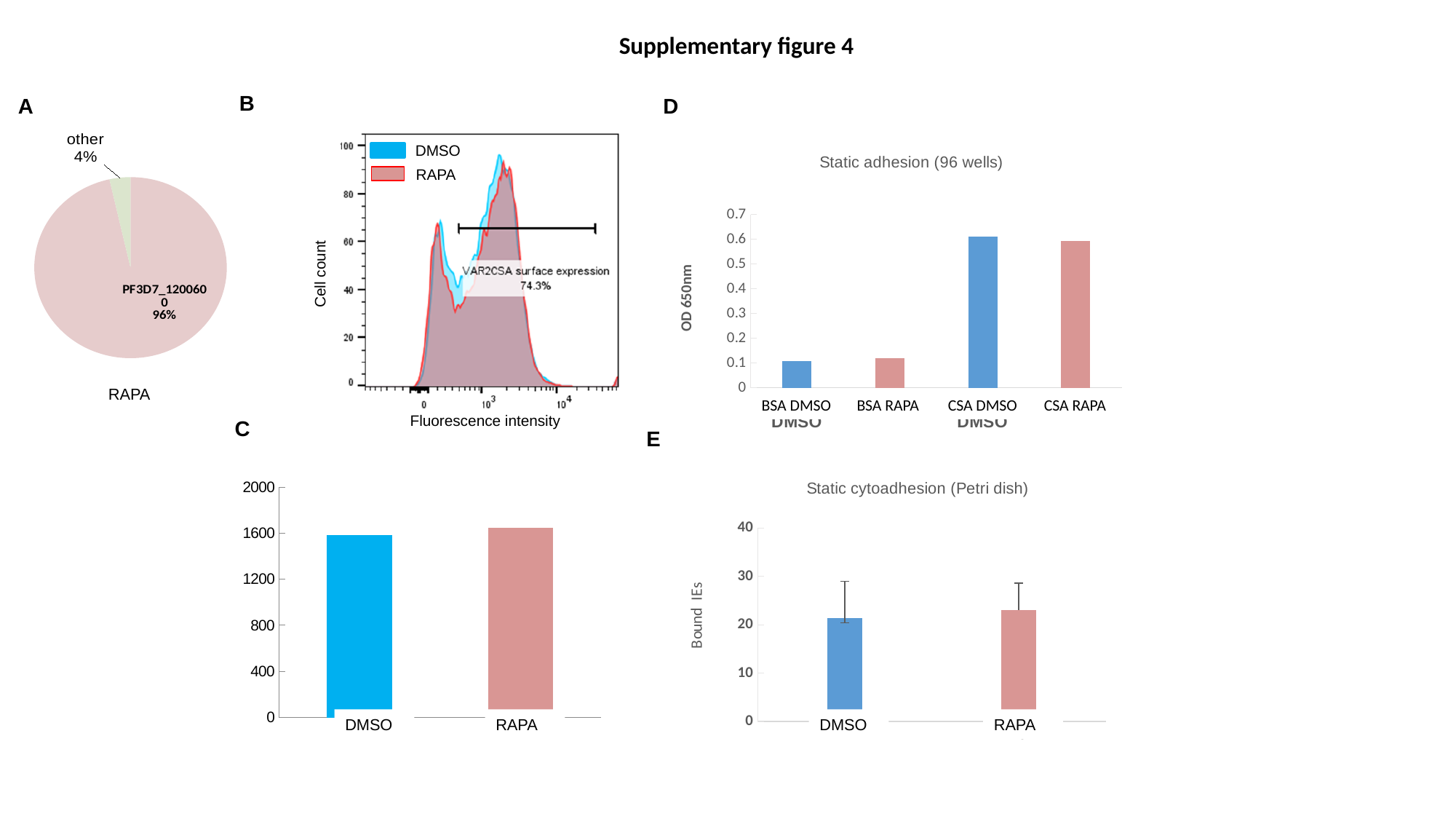

Supplementary figure 4
B
A
D
#### Chart
| Category | NF54 EPCR 4X |
|---|---|
| PFD1235w/MAL7P1.1 | None |
| PF11_0521 | None |
| PF13_0003 | None |
| PF08_0141 | None |
| PFD0020c | None |
| PFA0015c/PFI1820w/MAL6P1.314 | None |
| PFA0015c | None |
| PF08_0140 | None |
| MAL6P1.316 | None |
| PFL0020w | None |
| MAL6P1.4 | None |
| PF11_0007 | None |
| PF08_0142 | None |
| PFE0005w | None |
| PFA0005w | None |
| PFA0765c | None |
| PFC1120c/PFC0005w | None |
| PFD0005w | None |
| PFI0005w | None |
| PF13_0364 | None |
| PF07_0139 | None |
| PFB1055c | None |
| PF10_0406 | None |
| PFL0005w | None |
| PFB0010w | None |
| PFL2665c | None |
| PF13_0001 | None |
| MAL6P1.1 | None |
| PFD1245c | None |
| PFI1830c | None |
| PF10_0001 | None |
| PFL0935c | None |
| PFL1955w/PFL1970w | None |
| PF08_0106 | None |
| MAL7P1.50 | None |
| PF08_0103 | None |
| MAL7P1.55 | None |
| PF07_0050 | None |
| PFL1950w | None |
| MAL6P1.252 | None |
| MAL7P1.56 | None |
| PFD0995c/PFD1000c | None |
| PFD0995c | None |
| PF07_0049 | None |
| PFD0630c/PFD0635c | None |
| PFD1005c/PFD1015c | None |
| PFD1015c | None |
| PFD0615c | None |
| PF07_0051 | None |
| PF07_0048 | None |
| PFL1960w | None |
| PFD0625c | None |
| PF11_0008 | None |
| PFI1820w | None |
| PFE1640w | None |
| PF3D7_1200600 | 96.47776462077836 |
| other | 3.522235379221641 |
DMSO
#### Chart: Static adhesion (96 wells)
| Category | |
|---|---|
| BSA DMSO | 0.1075 |
| BSA Rapa | 0.1205 |
| CSA DMSO | 0.6105 |
| CSA Rapa | 0.595 |
RAPA
Cell count
RAPA
BSA DMSO
BSA RAPA
CSA DMSO
CSA RAPA
Fluorescence intensity
C
E
#### Chart: Static cytoadhesion (Petri dish)
| Category | |
|---|---|
| CSA DMSO | 21.4 |
| Rapa | 23.0 |
#### Chart
| Category | |
|---|---|
| DMSO | 1581.0 |
| Rapa | 1648.0 |DMSO
RAPA
DMSO
RAPA

### Slide 5
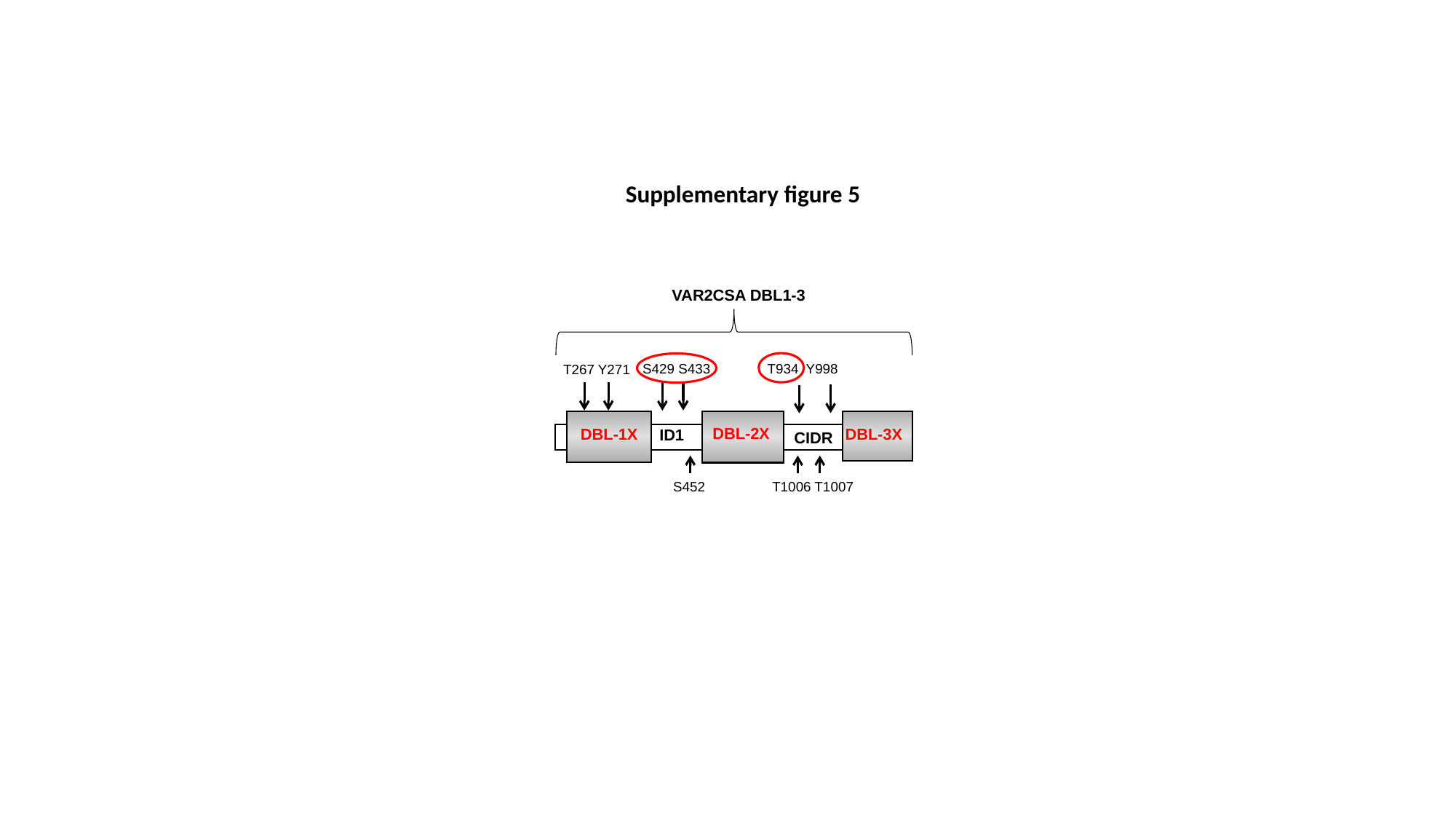

Supplementary figure 5
VAR2CSA DBL1-3
 S429 S433 T934 Y998
 T267 Y271
DBL-2X
DBL-1X
DBL-3X
IR1
ID1
CIDR
S452
 T1006 T1007
